# Impact of small RNAs in retrograde signalling pathways in *Arabidopsis thaliana*

**DOI:** 10.1101/798900

**Authors:** Kristin Habermann, Bhavika Tiwari, Maria Krantz, Stephan O. Adler, Edda Klipp, M. Asif Arif, Wolfgang Frank

## Abstract

Chloroplast perturbations activate retrograde signalling pathways causing dynamic changes of gene expression. Besides transcriptional control of gene expression different classes of small non-coding RNAs (sRNAs) act in gene expression control, but comprehensive analyses regarding their role in retrograde signalling is lacking. We performed sRNA profiling in response to norflurazon (NF) that provokes retrograde signals in *A. thaliana* wild type and the two retrograde signalling mutants *gun1* and *gun5*. The RNA samples were also used for mRNA and long non-coding RNA (lncRNA) profiling to link altered sRNA levels to changes of their cognate target RNAs. We identified 122 sRNAs from all known sRNA classes that were responsive to NF in wild type. Strikingly, 140 and 213 sRNAs were found to be differentially regulated in both mutants indicating a retrograde control of these sRNAs. Concomitant with the changes in sRNA expression we detected about 1500 differentially expressed mRNAs in the NF treated wild type and around 900 and 1400 mRNAs that were differentially regulated in the *gun1* and *gun5* mutant with a high proportion (~30%) of genes encoding plastid proteins. Furthermore, around 20% of predicted miRNA targets code for plastid localised proteins. The analyses of sRNA-target pairs identified pairs with an anticorrelated expression as well pairs showing other expressional relations pointing to a role of sRNAs in balancing transcriptional changes upon retrograde signals. Based on the comprehensive changes in sRNA expression we assume a considerable impact of sRNAs in retrograde-dependent transcriptional changes to adjust plastidic and nuclear gene expression.

**Significance statement:** Perturbations of plastid functions trigger retrograde signalling to adjust plastidic and nuclear gene expression, however, the role of small non-coding RNAs acting as regulators in these pathways is not well understood. We analysed small non-coding RNA expression in response to retrograde signals in *A. thaliana* wild type and two retrograde signalling mutants and identified members of all known small non-coding RNA classes pointing to a functional role of these RNA classes in retrograde pathways.

## Introduction

Both mitochondria and chloroplasts are characteristic organelles of eukaryotes and have evolved through the endosymbiosis of distinct prokaryotic progenitors, an evolutionary event that has led to the endosymbiotic theory (Goksoyr, 1967). Whereas the endosymbiosis of α-proteobacteria led to the formation of mitochondria in all common eukaryotes, plastids present in the green lineage of life have evolved from endosymbiotic cyanobacteria. However, both contained a considerable large genome encoding a few thousands of proteins. During the functional internalisation and transformation into plastids the majority of the cyanobacterial genome was transferred into the nuclear DNA of the host organism and only about 200 protein coding genes remained in the plastid genome. Consequently, most multiprotein complexes within the plastids are formed by organellar and nuclear encoded proteins requiring a well-coordinated gene expression of both genomes (Zimorski et al., 2014). In addition, all nuclear encoded proteins required for chloroplast development and functions have to be imported into the plastids (Martin et al., 2002). The coordination of gene expression in both compartments is controlled by two major pathways that balance nuclear and organellar gene expression during development and in response to environmental stimuli. The expression of organellar genes is controlled by nucleus-to-plastid signals known as anterograde control and, vice versa, nuclear gene expression is controlled by plastid-to-nucleus signals referred to as retrograde signalling (Kleine and Leister, 2016). Both pathways act together to maintain the cellular homeostasis and it is proposed that they are mediated by several factors. For example, changes in plastid gene expression affect nuclear gene expression since lincomycin that inhibits plastidal translation prevents the expression of nuclear encoded photosynthesis associated nuclear genes (*PhANGs*) (Sullivan and Gray, 1999) indicating that the plastid senses defective gene expression and generates retrograde signals to coordinate nuclear gene expression (Gray et al., 2003). Moreover, norflurazon (NF) that is a specific inhibitor of the enzyme phytoene desaturase producing β-carotenoids from phytoene was also shown to cause repression of *PhANGs* (Woodson et al., 2011). Carotenoids are part of the light-harvesting complexes and protect the cells from photooxidative damage (Kim and Apel, 2013). In the presence of NF the chloroplast suffers from photooxidation leading to characteristic bleaching symptoms of the green plant tissues caused by the degradation of chlorophyll (Breitenbach et al., 2001). Several decades ago *A. thaliana* mutant screens were performed to identify factors which specifically block the expression of *PhANGs* under conditions of chloroplast developmental prevention (Susek et al., 1993, Larkin et al., 2003, Mochizuki et al., 2001, Gray et al., 2003, Saini et al., 2011, Meskauskiene et al., 2001, Gutierrez-Nava Mde et al., 2004, Ball et al., 2004, Rossel et al., 2006). The promoter of the *LHCB* gene (light harvesting complex of photosystem II) was fused to the *GUS* reporter gene to monitor altered *LHCB* expression in ethyl methanesulfonate (EMS) mutagenised *A. thaliana* seedlings in response to NF (Susek et al., 1993). *A. thaliana* wild type plants grown in the presence of NF generate retrograde signals to inhibit the expression of *PhANGs* and the mutagenised seedlings were screened for de-repressed *LHCB* expression in response to NF indicative for a disturbed retrograde signal. Several GENOME UNCOUPLED (*gun*) mutants displaying a de-repressed *LHCB* expression were isolated and the affected genes were subsequently identified (Susek et al., 1993). Interestingly, five different *gun* mutants, *gun2* to *gun6*, are affected in the tetrapyrrole biosynthesis pathway (TPB). The mutation in the *gun5* mutant results in a defective regulatory CHLH subunit of the magnesium-chelatase (Mochizuki et al., 2001). The *gun4* mutation also affects the subunit of the magnesium-chelatase, leading to an increased efficiency. The *gun2*, *gun3* and *gun6* mutants are impaired in the heme oxygenase, phytochromobilin synthase and Fe-chelatase, respectively (Woodson et al., 2011, Woodson et al., 2013). Based on these studies it has been proposed that chloroplast metabolites may act as retrograde signals controlling the expression of *PhANGs* (Kakizaki et al., 2009). Another metabolite is 3′-phosphoadenosine 5′-phosphate (PAP) which acts as a mobile signal in the RNA metabolism since it blocks exoribonucleases controlling steady-state levels of nuclear encoded RNAs that are necessary for plastid development (Estavillo et al., 2011). The *gun1* mutant is not related to the remaining *gun* mutants, because it shows the same phenotype under NF as well as under lincomycin treatment. *GUN1* encodes a member of the chloroplast-localised pentatricopeptide repeat proteins (PPR) that usually act in posttranscriptional processes (Tadini et al., 2016). The *gun1* mutant is able to perceive signals from the tetrapyrrole biosynthesis pathway, plastid gene expression and redox state, but the mode of action of *GUN1* in retrograde signalling remains unknown (Kleine and Leister, 2016).

Beside these initial studies other screens were performed to identify further plastid-derived retrograde signals (Pogson et al., 2008) and the redox state (Mayfield and Taylor, 1984), intermediates of tetrapyrrole biosynthesis (Larkin et al., 2003), sugar and reactive oxygen species (ROS) (op den Camp et al., 2003) were suggested to be plastid-derived signals involved in retrograde signalling.

To study the impact of the individual retrograde signals on transcriptional changes several studies have been performed using a selection of available retrograde signalling mutants (Koussevitzky et al., 2007, Strand et al., 2003). Strand *et al*. (2003) compared the transcriptome of *A. thaliana* wild type and *gun1*, *gun2* and *gun5* mutants in response to a 6 day NF treatment by microarray studies (Strand et al., 2003). Another study (Koussevitzky et al., 2007) also analysed the transcriptome of *A. thaliana* wild type, *gun1* and *gun5* mutants by using microarrays after NF treatment for 5 days. In both studies it has been observed that *PhANGs* are predominantly downregulated in wild type whereas *PhANGs* become de-repressed in both *gun* mutants. Furthermore, these studies revealed a strong correlation between the *gun1* and the *gun5* mutant since a large number of genes were consistently regulated in both mutants.

To date, all studies analysing gene expression in various retrograde signalling mutants focused on the analysis of protein coding genes. However, it is well known that classes of non-coding RNAs (ncRNAs) including long ncRNAs as well as small ncRNAs have important functions in diverse biological processes since they mainly act in the control of gene expression. However, only a single study was reported addressing the potential role of small RNAs (sRNAs) in retrograde signalling (Fang et al., 2018).

This specific class of non-coding RNA comprises sRNAs with a size of 20-24 nucleotides that are able to regulate gene expression. They can interfere with nuclear transcription by regulating epigenetic modifications (Holoch and Moazed, 2015, Bannister and Kouzarides, 2011, Khraiwesh et al., 2010) or they act post-transcriptionally by targeting mRNAs mediating RNA cleavage or translational inhibition (Meister and Tuschl, 2004, Bartel, 2004, Kim, 2005). Small non-coding RNAs can be divided into two classes on the basis of their origin: hairpin RNA (hpRNA) and small interfering RNA (siRNA) (Axtell, 2013a). SiRNAs are processed from double-stranded RNA (dsRNA) precursors, whereas hpRNAs have single-stranded precursors that fold back into a characteristic hairpin structure. MicroRNAs (miRNAs) present one of the most important and well known class of hpRNA that mediate regulation of target gene expression (Meyers et al., 2008). MiRNAs are processed from characteristic stem-loop primary precursor transcripts that are encoded by *MIR* genes. The mature miRNAs are processed from the hairpin region of these precursors by DICER-LIKE1 (DCL1) enzymes (Park et al., 2002). Further, the miRNA is loaded to the ARGONAUTE1 (AGO1) containing RNA-induced silencing complex (RISC) that guides the miRNA to its target RNA by means of sequence complementarity and mediates subsequent target RNA cleavage or translational inhibition (Wierzbicki et al., 2008, Voinnet, 2009). Until now only one recent study reports on a functional role of miRNAs in retrograde signalling (Fang et al., 2018). In this study tocopheroles and PAP were identified to have an effect on miRNA biogenesis. It was shown that tocopherols positively regulate PAP accumulation which is an inhibitor of the exonuclease 2 (XRN2) (Fang et al., 2018) that negatively regulates mRNA and pri-miRNA levels by degradation of 5’ uncapped mRNA. Thus, increased PAP levels lead to an enhanced repression of XRN2 which in turn results in an increased pri-miRNA accumulation (Fang et al., 2018). Moreover, the generation of PAP is dependent on miR395 since it mediates cleavage of the mRNA encoding ATP sulfurylase (APS) the enzyme catalysing the initial step of PAP synthesis (Fang et al., 2018). Two other classes of sRNAs are formed from dsRNAs that derive from endogenous transcripts and generate natural antisense transcript-derived siRNA (nat-siRNA) or trans-acting siRNA (ta-siRNA) based on their specific biogenesis pathways. Nat-siRNAs are generated from two genes that encode overlapping transcripts in antisense orientation. Upon simultaneous transcription they produce complementary transcripts that are able to generate dsRNA molecules (Borsani et al., 2005). From the dsRNA DCL2 first processes a 24 nt nat-siRNA which then targets one of the initial overlapping transcripts and mediates its cleavage. Subsequently, RNA-dependent RNA polymerase 6 (RdR6) converts the cleavage products into dsRNA that serve as substrates for DCL1 to generate 21 nt nat-siRNAs that target one of the original overlapping transcripts mediating its cleavage and subsequent degradation (Borsani et al., 2005). According to their chromosomal arrangement and localisation *cis*- or *trans*-nat-siRNAs can be distinguished (Yuan et al., 2015). If two genes are located as overlapping genes on the opposite DNA strands in an identical genomic region and both genes are transcribed simultaneously they are able to generate *cis*-nat-siRNA. If two genes encoding transcripts with reverse complementary sequences are separated from each other at different genomic locations, they are also able to form dsRNAs upon simultaneous transcription and give rise to *trans*-nat-siRNA (Lapidot and Pilpel, 2006). In both cases the nat-siRNAs have the same function in targeting one of the initial transcripts irrespective of the different chromosomal arrangement. The first identified nat-siRNA was shown to have an important function in salt stress adaptation of *A. thaliana* (Borsani et al., 2005) where it is involved in the regulation of proline biosynthesis. Unlike nat-siRNAs, ta-siRNA generation is triggered by miRNAs since ta-siRNA precursor transcripts are cleaved in a miRNA-dependent manner. The cleaved *TAS* precursor transcript is further processed and converted into dsRNA by RDR6 and processed into phased 21 nt ta-siRNA duplexes by DCL4 proteins which are then loaded into a RISC complex to control target RNAs (Chen, 2009).

In addition to sRNAs that are processed from specific RNA precursors, long non-coding RNAs (lncRNAs) were shown to have important functions in the control of gene expression. These lncRNAs comprise RNA molecules with a size larger than 200 nucleotides that are transcribed by RNA Pol II, III, IV or V and do not possess protein coding capacities (Wierzbicki et al., 2008, Dinger et al., 2009). These lncRNAs have diverse roles and exert their function by various mechanisms. One specific role of lncRNAs is the regulation of mRNA splicing where they can either activate or inhibit specific splicing events (Ma et al., 2014). They also act in miRNA target mimicry, where the lncRNA harbours a miRNA binding site and thus mimics an mRNA target causing miRNA binding and sequestration. One well understood example for target mimicry in *A. thaliana* involves the lncRNA *IPS1* (INDUCED BY PHOSPHATE STARVATION1) that is specifically induced upon phosphate deficiency (Franco-Zorrilla et al., 2007) and harbours a miR399 binding site. MiR399 usually targets the *PHO2* mRNA encoding an ubiquitin-conjugating E2 enzyme which is able to tag proteins for degradation. Competitive binding of miR399 to *IPS1* causes elevated *PHO2* mRNA and protein levels which is required for survival under phosphate deficiency (Bari et al., 2006). In addition, lncRNAs can act as transcriptional regulators of gene expression by controlling epigenetic modifications. For example, the two lncRNAs COLD INDUCED LONG ANTISENSE INTRAGENIC RNA (*COOLAIR*) (Swiezewski et al., 2009) and COLD ASSISTED INTRONIC NONCODING RNA (*COLDAIR*) (Heo and Sung, 2011) can guide chromatin modifications at the FLOWERING LOCUS C (*FLC*) which inhibits flowering under low temperature by epigenetic repression of the *FLC* locus.

With the exception of a recently reported study (Fang et al., 2018), the role of sRNAs in retrograde signalling has not been analysed yet and information on the role of lncRNAs in retrograde control is completely lacking. To gain information whether these classes of non-coding RNA may act in retrograde signalling we made use of two well characterised mutants that are affected in plastid-to-nucleus signalling events. *A. thaliana gun1* and *gun5* mutants were grown under standard conditions as well as in the presence of NF and RNA expression profiles were compared to wild type controls. RNA profiles were monitored for mRNA, lncRNA and all sRNA classes to identify changes in RNA expression that depend on specific retrograde signalling pathways. Furthermore, sRNA profiles were correlated to mRNA and lncRNA profiles to identify putative functional sRNA-RNA targets which are modulated by or contribute to retrograde signals.

## Results

### De Novo sRNA sequencing after norflurazon treatment

To identify sRNAs that may act in retrograde signalling pathways, four day old seedlings of *A. thaliana* wild type (WT) as well as seedlings of the two retrograde signalling mutants *gun1* and *gun5* were treated for four days with 5 µM NF under continuous light (Figure 1a) and sRNA sequencing was performed from all samples. Each treatment was performed in six independent biological replicates and sRNA sequencing yielded a minimum of 5 million reads per replicate. The length distribution of all sRNA reads was analysed and we observed an enrichment of reads with a length of 21 and 24 nt (Figure 1b, c and d). The 21 nt peak corresponds to an expected enrichment of miRNAs, and ta-siRNAs and nat-siRNAs whereas the 24 nt peak complies with enriched repeat-associated sRNAs. The ShortStack sRNA analysis software has been used to map the sRNA data set against different reference databases including miRNAs (miRBase), lncRNAs (Araport 11), phasiRNAs (Howell et al., 2007) and previously predicted NATs-pairs (Zhang et al., 2012, Yuan et al., 2015, Jin et al., 2008) (Table S1) and to perform subsequent differential expression analysis of mapped sRNAs (Table S2).

**Figure 1.**
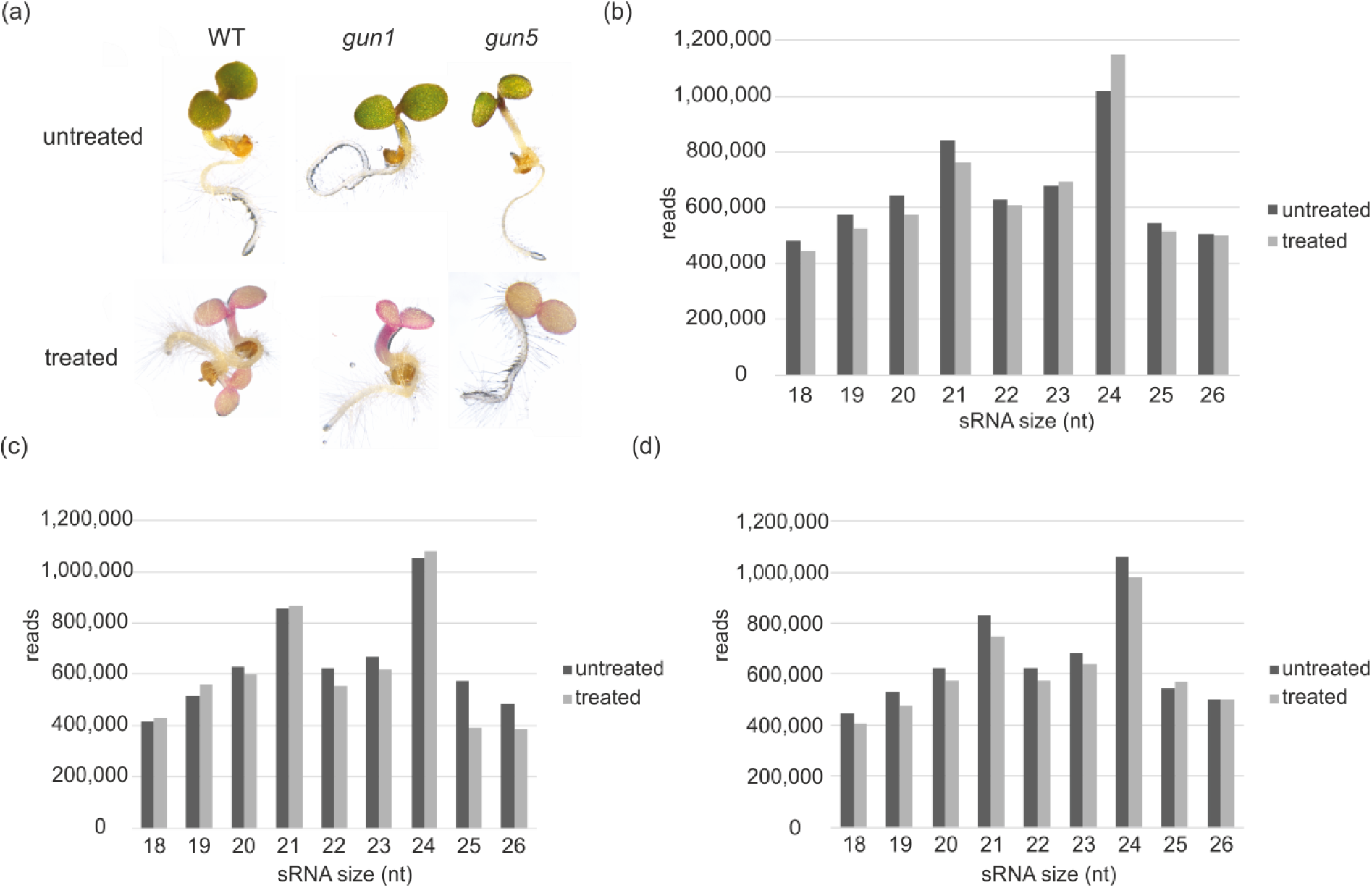
Effect of NF on the plants and sRNA size distribution. (a) Representative images of seedlings from WT, *gun1* and *gun5* mutants grown for four days in the presence or absence of NF. Size distribution of total sRNA reads (average of six biological replicates per sample) obtained from untreated and NF treated WT (b), *gun1* (c) and *gun5* (d) mutants.

DeSeq2 was used to calculate the differentially expression of sRNAs between the samples with a special focus on sRNAs that were differentially expressed in NF treated samples with respect to their untreated controls, and sRNAs that were differentially expressed in NF treated *gun* mutants compared to the NF treated wild type. All sRNAs with an at least 2 fold up- or downregulation and a q-value of ≤ 0.05 were considered as differentially expressed sRNAs. Specific sRNA clusters arising from different ncRNA classes were found to be differentially expressed (Figure 2). These classes include mature miRNAs, *cis*-NATs-pairs and *trans*-NATs-pairs as well as sRNAs derived from lncRNAs. Upon growth on normal media without NF we identified only a small number of differentially regulated sRNAs in the *gun* mutants as compared to the wild type whereas the number of differentially regulated sRNAs between the mutants and wild type strongly increased upon NF treatment. Comparing the untreated wild type and the untreated *gun* mutants we identified only 6 miRNAs, and sRNA clusters arising from 11 lncRNAs, one phasiRNA, 12 *cis*-NATs-pairs and 73 *trans*-NATs-pairs to be differentially expressed in the *gun1* mutant. In the *gun5* mutant five miRNAs and sRNA clusters arising from 18 lncRNAs, 12 *cis*-NATs-pairs and 27 *trans*-NATs-pairs were differentially regulated as compared to the control (Table S3).

**Figure 2.**
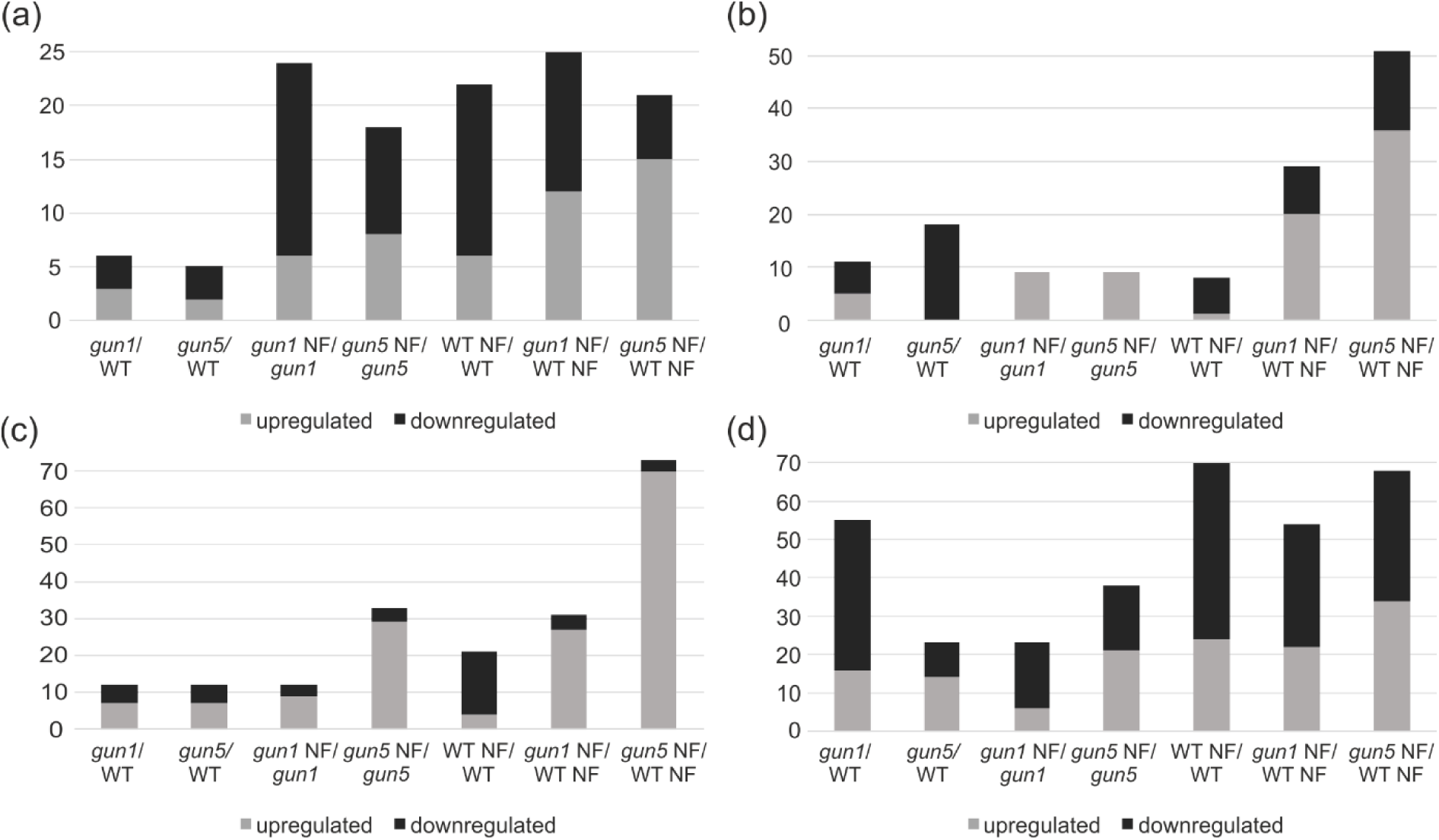
Differentially expressed sRNAs within the different samples. Overview of differentially regulated sRNAs between the different samples (FC ≤-2 and ≥+2; FDR ≤ 0.05) subdivided into specific sRNA classes. miRNAs (a), sRNAs deriving from lncRNA (b), *cis*-NATs-pairs (c) and *trans*-NATs-pairs (d). The up- and downregulation of the members of each class is depicted by grey (up) and black (down) partitions of the respective bars.

NF treatment caused an increase in the number of differentially expressed sRNAs (Table S3) in each genotype and provoked remarkable differences between wild type and both *gun* mutants indicating a considerable sRNA regulation by retrograde signals. After NF treatment in the wild type (WT NF) we detected 22 miRNAs (belonging to 15 miRNA families) and sRNA clusters arising from eight lncRNAs, one phasiRNA, 22 *cis*-NATs-pairs and 68 *trans*-NATs-pairs to be differentially expressed compared to the untreated control. Beside this, sRNA expression in response to NF in both mutants were compared to the respective untreated controls. 24 differentially regulated miRNAs (17 different miRNA families) were detected in the treated *gun1* mutant (*gun1* NF). Along with these miRNAs nine lncRNAs, one phasiRNA, 12 *cis*-NATs-pairs and 32 *trans*-NATs-pairs were found to produce differentially expressed sRNAs in *gun1* NF compared to the untreated *gun1* control. In the NF treated *gun5* mutant (*gun5* NF) 18 miRNAs (14 different miRNA families) and sRNA clusters arising from nine lncRNAs, 33 *cis*-NATs-pairs and 41 *trans*-NATs-pairs were found to be differentially regulated as compared to the untreated *gun5*. Strikingly, we detected 25 miRNAs (belonging to 14 different miRNA families) and sRNA clusters arising from 32 lncRNAs, one phasiRNA, 35 *cis*-NATs-pairs and 117 *trans*-NATs-pairs to be differentially expressed in *gun1* NF compared to the NF treated wild type. We observed a similar high number of differentially regulated sRNAs in the NF treated *gun5* mutant compared to NF treated wild type, namely 21 miRNAs (belonging to 17 different miRNA families) and sRNA clusters arising from 51 lncRNAs, 74 *cis*-NATs-pairs and 167 *trans*-NATs-pairs. Thus, the high number of differentially expressed sRNAs detected in both *gun* mutants in response to NF implies a strong regulation of sRNAs that underlies specific retrograde signals in these mutants. Most of the changes affect changes in miRNA and nat-siRNA expression and we mainly focus on these sRNA classes with regard to their differential expression and further target analysis to predict the regulatory functions of these sRNAs (Figure 2, Table S3).

### Analysis of differentially expressed miRNAs

Since beside a recent analysis of tocopherol-responsive miRNAs (Fang et al., 2018) nothing is known about the role of miRNAs in retrograde signalling we analysed changes in miRNA expression in response to NF in *A. thaliana* wild type as well as in the *gun1* and *gun5* retrograde signalling mutants. The comparison of differentially expressed miRNAs between the samples is shown in a hierarchically clustered (UPGMA) heatmap (Figure 3a). Only a low number of differentially expressed miRNAs was observed in the untreated mutants compared to the wild type control. In *gun1* only six differentially expressed miRNAs were detected (three upregulated and three downregulated) and only two upregulated and three downregulated miRNAs were detected in the untreated *gun5* mutant compared to the wild type control.

**Figure 3.**
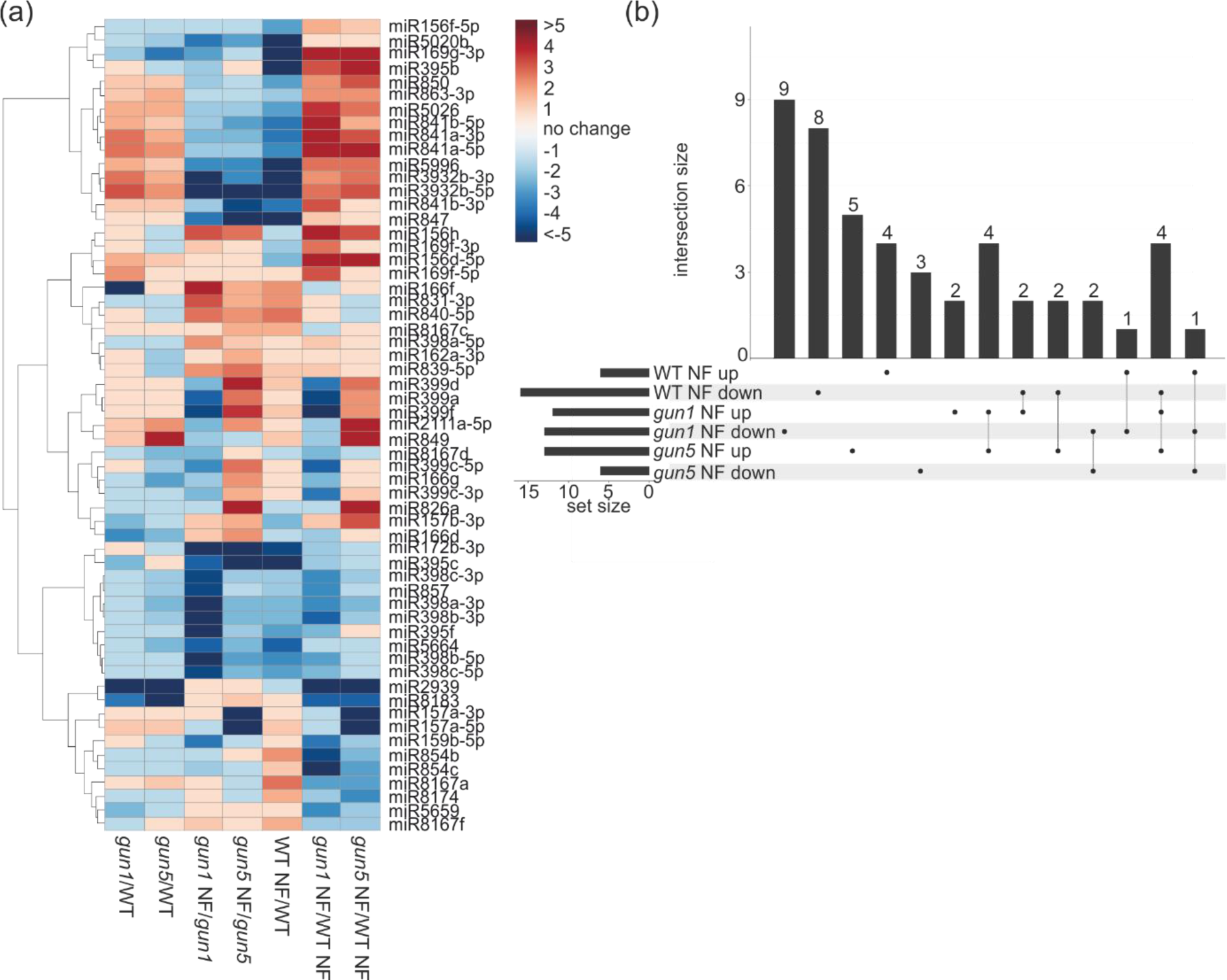
Behaviour of the differentially expressed miRNAs. (a) Hierarchically clustered (UPGMA) heatmap depicting miRNAs which are differentially regulated in at least one sample. (b) Upset plot depicting the number of differentially expressed miRNAs in response to NF in WT (WT NF/WT) and both *gun* mutants (*gun1* NF/WT NF and *gun5* NF/WT NF).

We hypothesised that miRNAs play a role in retrograde signalling which should be reflected by an enrichment of differentially expressed miRNAs after NF treatment. In our sequencing data we observe a remarkable increase in the number of differentially expressed miRNAs in response to NF treatment with a similar number of NF-responsive miRNAs in the three analysed genotypes (Figure 3b). In total, we observe 22 miRNAs to be differentially regulated in the NF treated wild type compared to the untreated control. Out of these only six miRNAs are upregulated and 16 miRNAs are downregulated. 24 miRNAs are differentially expressed in the NF treated *gun1* compared to the untreated *gun1* mutant (six upregulated and 18 downregulated) and eight miRNAs are upregulated and 10 downregulated in the NF treated *gun5* mutant compared to the untreated *gun5* control.

Interestingly, we further detected miRNAs that seem to be controlled by retrograde signals as de-repressed miRNAs were observed in the *gun1* and *gun5* mutants in response to NF treatment which is reminiscent to the de-repression of *PhANGs* in these mutants. We focused on miRNAs with altered expression levels in the treated wild type compared to the control condition and correlated them with differentially expressed miRNAs in the NF treated *gun* mutants (Figure 3b). In response to NF two miRNAs (miR169g-3p and miR5996) show the pattern of de-repression in both *gun* mutants similar to de-repressed *PhANGs* and one miRNA (miR3932-5p) was de-repressed only in the NF treated *gun5* mutant. Furthermore, five miRNAs were downregulated in the treated wild type compared to the control and were upregulated in at least one NF treated *gun* mutants (*gun* NF/WT NF). We also identified two miRNAs which seemed to be controlled by retrograde signals in an opposite manner. These two miRNAs were found to be upregulated in the treated wild type compared to the control and downregulated in at least one of the treated *gun* mutants (*gun* NF/WT NF). In addition, we found miRNAs which showed a specific regulation restricted to NF treated *gun* mutants when compared to the treated wild type. Two miRNAs were found to be downregulated in both treated *gun* mutants as compared to the NF treated wild type. Moreover, nine miRNAs were specifically downregulated in the NF treated *gun1* mutant and the expression of three miRNAs was reduced in the treated *gun5* mutant compared to the treated wild type. In addition, two miRNAs were upregulated in the treated *gun1* mutant and five miRNAs were upregulated in the treated *gun5* mutant compared to the treated wild type. We could also detect four upregulated miRNAs common for both treated *gun* mutants compared to the treated wild type.

### Differentially regulated nat-siRNAs

In *A. thaliana*, nat-siRNAs are processed in subsequent steps by DCL1 and DCL2 from overlapping transcripts. They target one of the transcripts by reverse sequence complementarity and mediate its cleavage. To identify nat-siRNA from our sRNA sequencing data, we made use of different accessible databases that comprise experimentally validated and computationally predicted NATs-pairs (Zhang et al., 2012, Yuan et al., 2015, Jin et al., 2008). According to their genomic location NATs-pairs can be distinguished into *cis*- and *trans*-NATs-pairs. *Cis*-NATs-pairs are generated from two overlapping genes located on opposing DNA strands within an identical genomic region whereas *trans*-nat-siRNAs are generated from transcripts that are encoded from separated genomic regions. In total, the reference databases used for the mapping (Table S1) of the sequenced sRNAs encompass 1902 *cis*-NATs-pairs and 2171 *trans*-NATs-pairs that were analysed with ShortStack to detect differentially expressed nat-siRNA clusters.

In the untreated *gun1* and *gun5* mutants as compared to the wild type, we identified six *cis*-NATs-pairs producing differentially regulated 21 nt nat-siRNA clusters that were commonly regulated in both mutants and six pairs with distinct regulations in each of the mutants (Figure 2c). Beside this, 57 and 23 *trans*-NATs-pairs were detected to produce differentially regulated nat-siRNA in the *gun1* and *gun5* mutant, respectively (Figure 2d).

Upon NF treatment we detected 21 *cis*-NATs-pairs (Figure 4a) and 70 *trans*-NATs-pairs (Figure 4b) in the wild type compared to the untreated control that produce differentially regulated 21 nt nat-siRNA clusters from at least one transcript of these NATs pairs. Most of these nat-siRNAs were found to be downregulated with 17 and 46 loci from *cis*-NATs-pairs and *trans*-NATs-pairs, respectively. In the treated *gun1* mutant nat-siRNAs from 12 *cis*-NATs-pairs were detected to be differentially expressed and most of them were upregulated compared to the untreated *gun1* mutant (Figure 2c). Moreover, 23 differentially regulated *trans*-NATs-pairs producing 21 nt nat-siRNA clusters were identified to be differentially regulated in the treated *gun1* mutant compared to the untreated *gun1* control and most of them were downregulated. In the NF treated *gun5* mutant compared to the untreated *gun5* control we identified 33 *cis*-NATs and 38 *trans*-NATs generating differentially expressed nat-siRNAs. Here, we observed a higher number of upregulated nat-siRNA with 29 *cis*-NATs-pairs as well as 21 *trans*-NATs-pairs (Figure 2c and d).

**Figure 4.**
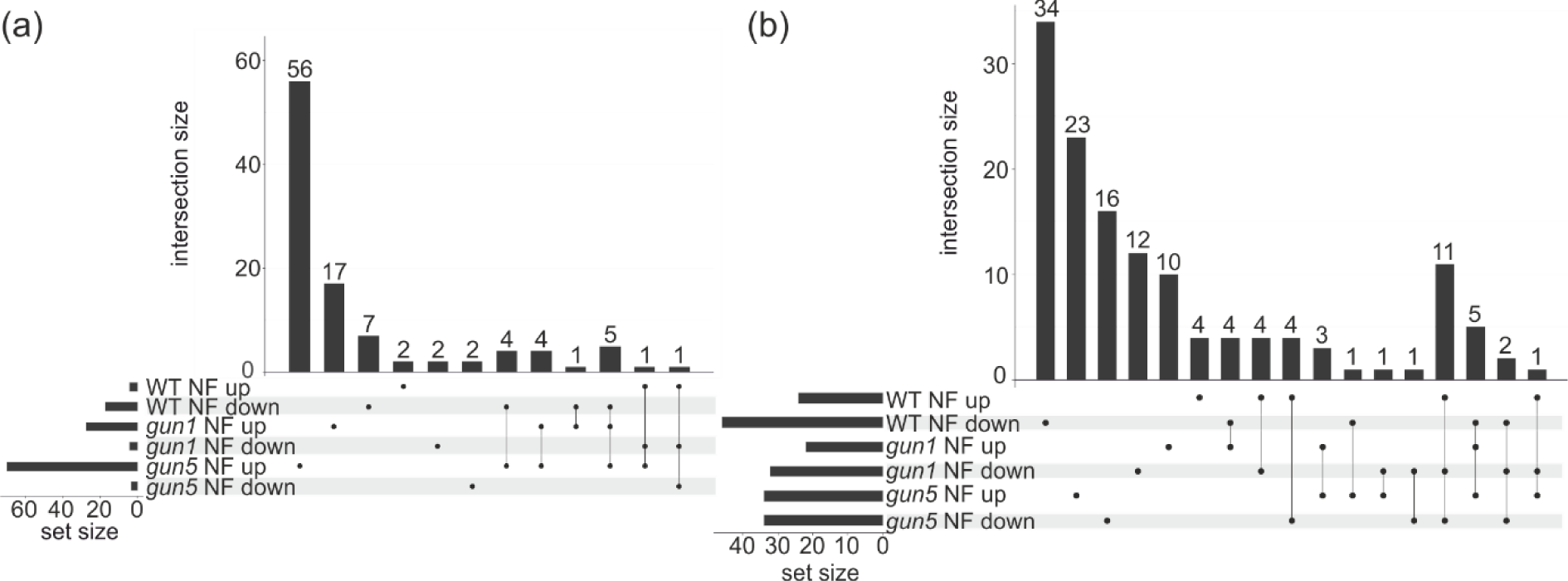
Distribution of differentially expressed *cis*- and *trans*-nat-siRNA in response to NF. UpSet plot showing differentially regulated nat-siRNA in response to NF treatment in *A. thaliana* WT and the *gun1* and *gun5* mutants. *cis*-NATs-clusters (a) and *trans*-NATs-clusters (b) in response to NF in WT (WT NF/WT) and both *gun* mutants (*gun1* NF/WT NF and *gun5* NF/WT NF).

The overlap and co-regulation as well as specific expression of the differentially expressed *cis*-NATs-pairs and *trans*-NATs-pairs that produce differentially regulated 21 nt nat-siRNA clusters were analysed between the samples and are shown in an UpSet plot (Figure 4). We focused on the analysis of NF-responsive differentially expressed nat-siRNAs in the wild type that should provide information on nat-siRNAs that are controlled by retrograde signals. Moreover, we compared NF treated wild type with both NF treated *gun* mutants to identify NF-responsive nat-siRNA misregulation that is caused by the perturbed retrograde signals in these mutants. We detected 31 *cis*-NATs-pairs and 54 *trans*-NATs-pairs that produce differentially expressed nat-siRNAs in the NF treated *gun1* mutant compared to the treated wild type. In the NF treated *gun5* mutant we detected 73 *cis*-NATs-pairs and 68 *trans*-NATs-pairs that produce differentially regulated 21 nt nat-siRNA clusters compared to the treated wild type. For both *gun* mutants the majority of *cis*-derived nat-siRNAs were upregulated whereas the majority of *trans*-derived nat-siRNAs were downregulated in these mutants (Figure 4). We identified five *cis*-NATs-pairs generating nat-siRNAs that were downregulated in the treated wild type (WT NF/WT) and were upregulated in both treated *gun* mutants (*gun* NF/WT NF) thus representing the *gun*-specific de-repression of nuclear encoded *PhANGs* (Figure 4a). We also detected one nat-siRNA cluster produced from a *cis*-NATs-pair that displayed an opposing expression pattern and was upregulated in the treated wild type (WT NF/WT) and downregulated in both mutants (*gun* NF/WT NF). Within *trans*-derived nat-siRNAs we identified 19 sRNA clusters that were differentially regulated in response to NF in wild type compared to the untreated control and showed further differential regulation in response to NF in both *gun* mutants compared to the NF treated wild type (Figure 4b). Five of them resemble the *gun*-specific de-repression since they were downregulated in the wild type (WT NF/WT) and upregulated in both mutants (*gun* NF/WT NF). 11 *trans*-derived nat-siRNA displayed an opposite expression and were upregulated in the treated wild type (WT NF/WT) and downregulated in both treated *gun* mutants (*gun* NF/WT NF). In addition, two differentially expressed nat-siRNA from *trans*-NATs-pairs were downregulated within all three samples and another one was upregulated in the treated wild type (WT NF/WT) as well as in the treated *gun5* mutant (*gun5* NF/WT NF) and downregulated in the *gun1* (*gun1* NF/WT NF).

### Differentially regulated other sRNA classes

Besides the differentially expressed miRNAs and NATs-pairs we also found differentially expressed sRNA that are produced from lncRNAs and *PHAS* precursors (Figure 5 and Table S3).

**Figure 5.**
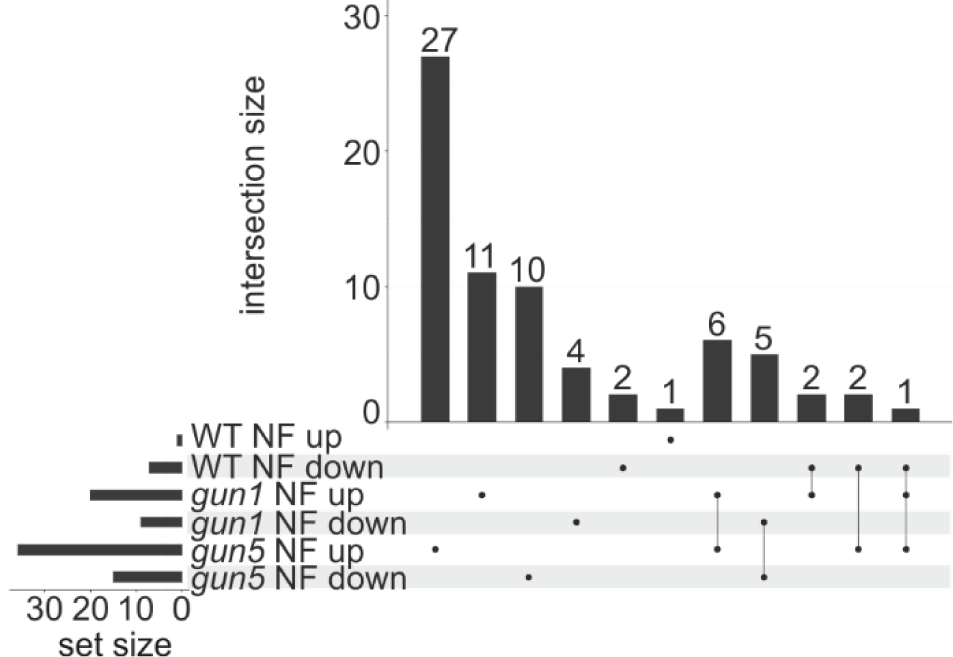
Distribution of differentially expressed sRNAs derived from lncRNAs in response to NF. UpSet plot depicting differentially regulated sRNAs derived from lncRNAs in response to NF in WT (WT NF/WT) and both *gun* mutants (*gun1* NF/WT NF and *gun5* NF/WT NF).

Similar to miRNAs and nat-siRNAs we detected only a small number of differentially regulated sRNA clusters derived from lncRNA precursors in the untreated genotypes and most of these sRNA clusters were downregulated in the untreated mutants compared to the wild type. In total, six out of 11 differentially expressed sRNA clusters produced from lncRNAs are downregulated in the *gun1* mutant and all 18 differentially expressed sRNA clusters in the untreated *gun5* mutant are downregulated compared to the wild type control.

When comparing the individual genotypes with and without NF treatment we also identified only a considerable small number of differentially expressed sRNAs. In the treated wild type eight sRNA clusters processed from lncRNA precursors were identified to be differentially expressed and seven of them were downregulated compared to the untreated control. For both treated *gun* mutants, we observed nine different upregulated sRNA clusters compared to their untreated control.

Comparing the NF treated *gun* mutants with the NF treated wild type we noticed an increase in the number of differentially expressed sRNAs (Figure 5). Generally, we observed a higher number of upregulated sRNA clusters produced from lncRNA precursors in both treated *gun* mutants compared to the treated wild type. 20 out of 29 differentially expressed sRNA clusters were detected to be upregulated in the NF treated *gun1* mutant compared to the treated wild type. For the treated *gun5* mutant 36 out of 51 differentially expressed sRNA clusters were found to be upregulated compared to the treated wild type. We detected only one sRNA cluster derived from an lncRNA precursor that is downregulated in the treated wild type compared to the untreated control and upregulated in both NF treated *gun* mutants. Two sRNA clusters are downregulated in the treated wild type compared to the control and upregulated in the treated *gun1* mutant and another two sRNA clusters derived from lncRNA precursors are downregulated in the treated wild type compared to the untreated control and upregulated in the treated *gun5* mutant. Furthermore, 11 sRNA clusters produced from lncRNA precursors are commonly regulated in both treated *gun* mutants with six of them upregulated and five downregulated compared to the treated wild type (Figure 5).

In addition to the lncRNA-derived sRNA clusters we identified two differentially expressed phasiRNAs. One phasiRNA derived from locus AT1G63070 is 5.8 fold upregulated in the untreated *gun1* mutant compared to the wild type control as well as 4.1 fold upregulated in the NF treated *gun1* mutant compared to the treated wild type. The second phasiRNA is produced from the locus AT5G38850 and is 2.6 fold downregulated in the treated wild type compared to the control and 3.5 fold downregulated in the treated *gun1* mutant compared to the untreated *gun1* mutant.

### Analysis of lncRNA and mRNA in the *gun* mutants

Until now only microarray analyses have been performed to characterise the de-repression of *PhANGs* in response to NF in *gun* mutants. Furthermore, only the expression of protein-coding mRNAs has been addressed whereas information on the NF-dependent regulation of lncRNAs is completely lacking (Koussevitzky et al., 2007, Strand et al., 2003). Therefore, we performed RNA sequencing from wild type, *gun1* and *gun5* mutants grown in the presence or absence of NF using the identical RNA samples that have been used for the sequencing of sRNAs described above to be able to perform subsequent studies on putative sRNA-mRNA/lncRNA relationships. RNA sequencing was performed from ribosomal depleted RNA including mRNA as well as lncRNA transcripts.

Three biological replicates of each sample were sequenced with paired-end reads obtaining an average number of approximately 12 million reads per replicate.

Using Tophat (Kim et al., 2013) the paired-end reads from each replicate were mapped against the *A. thaliana* genome deposited in Araport11 (Table S4) and differential expression of mRNA and lncRNA between the samples (Table S5 and S6) was calculated with Cuffdiff (Trapnell et al., 2010).

In total, 222 RNAs (212 mRNAs and 10 different classes of ncRNAs) were differentially regulated in the untreated *gun1* mutant compared to the untreated wild type. Out of these 92 RNAs were upregulated and 130 RNAs were downregulated (*gun1*/WT). In the untreated *gun5* (*gun5*/WT) we identified 175 differentially expressed transcripts (165 mRNAs and 10 ncRNAs) with 60 upregulated and 115 downregulated RNAs. In response to NF treatment the number of differentially expressed transcripts increased drastically and the major changes could be observed in the NF treated wild type (WT NF/WT). In total 1591 RNAs (1557 mRNAs and 34 ncRNAs) were differentially expressed in the treated wild type compared to the control including 435 upregulated and 1156 downregulated RNAs (WT NF/WT). Also high numbers of differentially expressed RNAs were observed in the NF treated mutants compared to their respective untreated controls. 1393 differentially expressed transcripts (1361 mRNAs and 32 ncRNAs) were identified in the treated *gun1* mutant compared to the untreated *gun1* including 511 upregulated and 882 downregulated RNAs (*gun1* NF/*gun1*). 1247 transcripts (1177 mRNAs and 70 ncRNAs) were differentially regulated in the treated *gun5* mutant compared to the untreated control (*gun5* NF/*gun5*) with 577 upregulated and 670 downregulated transcripts. Both the *gun1* and *gun5* mutant also display a high number of significantly changed transcripts after NF treatment compared to the treated wild type. 925 transcripts (905 mRNAs and 20 ncRNAs) are differentially regulated in the treated *gun1* mutant (*gun1* NF/WT NF) with about two third of the genes (595) upregulated and 330 genes downregulated. The same tendency was observed in the NF treated *gun5* mutant (*gun5* NF/WT NF) where 1364 differentially regulated RNAs (1319 mRNAs and 45 ncRNAs) were detected. Out of these 1019 RNAs showed an increased expression and 345 RNAs were downregulated. Representative transcripts belonging to different RNA classes and showing differential expression levels in the RNA sequencing data were selected for expression analyses by qRT-PCR to confirm the mRNA and lncRNA sequencing data (Figure S1). We selected various genes which were detected to be differentially expressed in the treated wild type compared to the untreated control as well as genes that were differentially regulated in both NF treated *gun* mutants compared to the treated wild type. Furthermore, we selected transcripts with varying abundance with two genes each displaying low, moderate and high abundance, respectively. In addition, we included expression analysis for one lncRNA that was found to be differentially regulated in all three samples.

### Classification of differentially expressed ncRNAs detected via ribosomal depleted RNA sequencing

NcRNAs encompass different types of RNAs, which can exert regulatory functions without coding for proteins and no studies have been reported that analyse the contribution of ncRNAs in the regulation of genes that underlie retrograde control. Besides mRNAs we also identified differentially expressed transcripts belonging to distinct ncRNA classes in our mRNA/lncRNA sequencing data that might play regulatory roles in retrograde signalling (Table S5 and S6). These different classes include lncRNA that may act in regulatory processes of gene expression as well as tRNA, rRNA and snoRNA that act in protein translation and splicing and usually have little regulatory functions. Under normal growth conditions we identified 10 differentially expressed ncRNAs in each of the *gun* mutants as compared to the untreated wild type (Table 1 and Figure 6a). The number of differentially expressed ncRNAs increased upon NF treatment indicating potential roles upon plastid perturbations that trigger retrograde signalling. In total we identified 34 differentially expressed ncRNAs in the NF treated wild type compared to the untreated control (Table 1). 11 of these were downregulated and 23 RNAs were upregulated (WT NF/WT). In the NF treated *gun1* and *gun5* mutants compared to their respective untreated controls we identified 32 and 70 differentially expressed ncRNAs, respectively. Interestingly, in the NF treated *gun* mutants as compared to treated wild type we observed 20 and 45 differentially expressed ncRNAs in *gun1* (*gun1* NF/WT NF) and *gun5* (*gun5* NF/WT NF), respectively.

**Figure 6.**
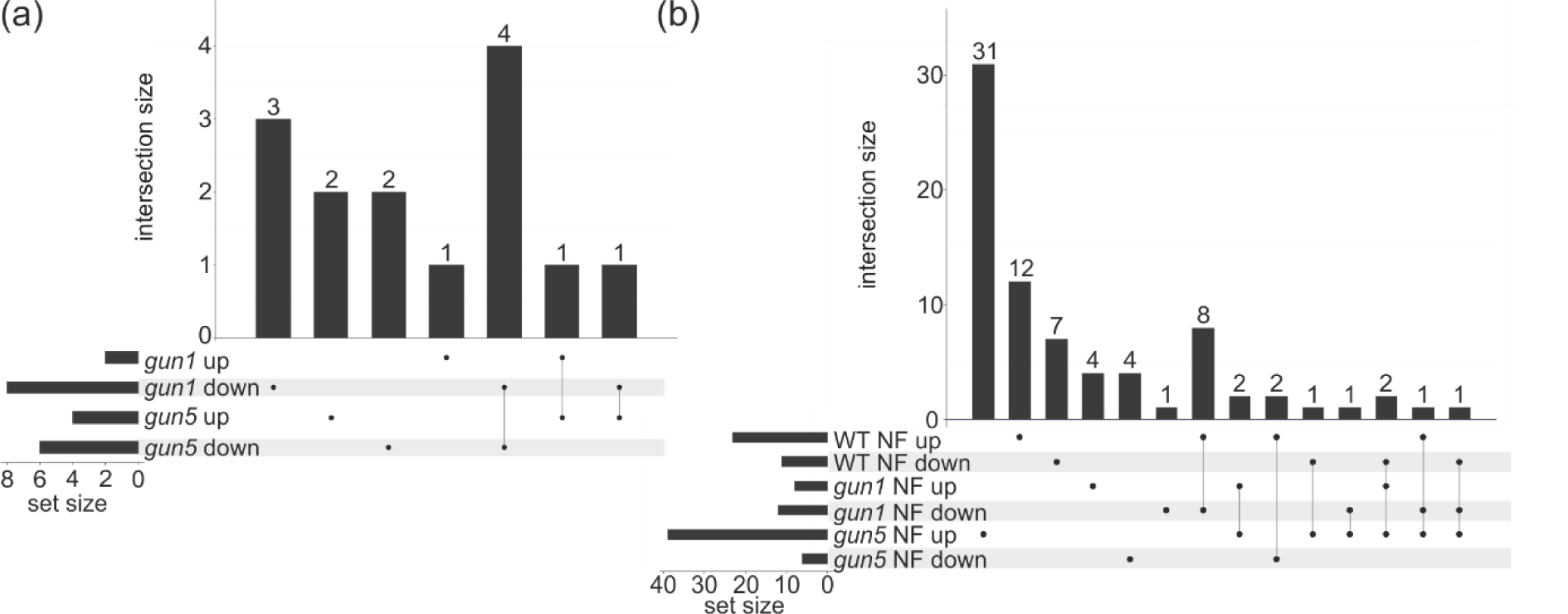
Distribution of differentially expressed ncRNAs in the untreated and treated samples. (a) Upset plot showing the distribution of differentially regulated ncRNAs in the untreated *gun* mutants compared to the untreated WT. (b) Upset plot depicting the distribution of differentially regulated ncRNAs in response to NF in WT (WT NF/WT) and both *gun* mutants (*gun1* NF/WT NF and *gun5* NF/WT NF).

**Table 1.** Overview of differentially expressed ncRNAs in response to NF in *A. thaliana* wild type, *gun1* and *gun5* mutants.

|  | ***gun1*/**<br>**WT** | ***gun5*/**<br>**WT** | **WT NF/**<br>**WT** | ***gun1* NF/**<br>***gun1*** | ***gun5* NF/**<br>***gun5*** | ***gun1* NF/**<br>**WT NF** | ***gun5* NF/**<br>**WT NF** |
| --- | --- | --- | --- | --- | --- | --- | --- |
| **lncRNA** | 6 | 5 | 15 | 13 | 34 | 11 | 20 |
| **snRNA** | 0 | 0 | 3 | 0 | 8 | 1 | 5 |
| **snoRNA** | 0 | 0 | 3 | 3 | 11 | 2 | 4 |
| **rRNA** | 0 | 0 | 2 | 0 | 1 | 1 | 0 |
| **tRNA** | 0 | 0 | 3 | 4 | 1 | 2 | 1 |
| **pseudogene** | 4 | 5 | 5 | 6 | 9 | 2 | 10 |
| **transcript**<br>**region** | 0 | 0 | 2 | 5 | 3 | 1 | 4 |
| **MIR precursor** | 0 | 0 | 1 | 1 | 2 | 0 | 0 |
| **antisense RNA** | 0 | 0 | 0 | 0 | 1 | 0 | 1 |
| **Total** | 10 | 10 | 34 | 32 | 70 | 20 | 45 |

An Upset plot (Figure 6b) was created to provide an overview of the distribution of differentially expressed ncRNAs between various samples (WT NF/WT, *gun1* NF/WT NF and *gun5* NF/WT NF). We identified two interesting lncRNAs (AT1G05562, AT4G13495) which represent the classical *gun*-related expression as these show a downregulation in response to NF treatment in wild type, but are upregulated in both NF treated *gun* mutants. Furthermore, three lncRNAs (AT3G01835, AT5G07325 and AT5G07745) were identified to be upregulated in the NF treated wild type (WT NF/WT) and downregulated in the treated *gun1* mutant (*gun1* NF/WT NF).

Another interesting lncRNA (AT4G13495) was de-repressed in both NF treated *gun* mutants compared to the treated wild type with 7.2 fold and 3.4 fold upregulation in *gun1* (*gun1* NF/WT NF) and *gun5* (*gun5* NF/WT NF), respectively, whereas this lncRNA was highly downregulated (10.5 fold) in the treated wild type. From our sRNA data we already detected sRNAs arising from this lncRNA and in agreement with the expression level of this lncRNA the total sRNAs generated from this transcript were downregulated in the treated wild type (−2.8 FC; WT NF/WT) and 3.4 fold upregulated in the treated *gun1* (*gun1* NF/WT NF). Interestingly, this lncRNA overlaps in sense direction with three individual miRNA precursors (MIR5026, MIR850 and MIR863) suggesting that these miRNAs can be processed from the individual precursors as well as from the overlapping lncRNA. In line with this hypothesis we observed a consistent differential expression of the lncRNA and the three individual miRNAs within the analysed samples (Table 2).

**Table 2.** Expression data for the lncRNA AT4G13495 that overlaps in sense with the individual miRNA precursors MIR5026, MIR850 and MIR863.

| ID | FC WT<br>NF/WT | q-value | FC *gun1*<br>NF/WT NF | q-value | FC *gun5*<br>NF/WT NF | q-value |
| --- | --- | --- | --- | --- | --- | --- |
| **AT4G13495** | -10.54 | 0.001 | 7.17 | 0.001 | 3.36 | 0.001 |
| **miR5026** | -2.63 | 0.085 | 4.08 | 0.004 | 3.08 | 0.042 |
| **miR850** | -2.58 | 0.077 | 2.61 | 0.068 | 3.81 | 0.007 |
| **miR863-5p** | -1.7 | 0.561 | 1.29 | 0.941 | 2.34 | 0.467 |
| **miR863-3p** | -1.51 | 0.538 | 2.7 | 0.025 | 2.61 | 0.045 |

In addition, we also identified two differentially regulated lncRNAs that overlap with mRNA transcripts in antisense and may act as precursors for the generation of nat-siRNAs. The first lncRNA transcript (AT3G51075) was found to be upregulated in the treated wild type compared to the control (FC of 2.3 WT NF/WT) as well as downregulated in the treated *gun1* compared to the treated wild type with a FC of −2.2 (*gun1* NF/WT NF). This lncRNA transcript overlaps with an mRNA transcript (AT3G51070) coding for an S-adenosyl-L-methionine-dependent methyltransferase. However, the expression of the overlapping mRNA transcript remained unchanged and we did not detect cognate nat-siRNAs processed from the overlapping region in our sRNA data. The second lncRNA that may act as a natural antisense transcript (AT1G05562) was differentially regulated in both NF treated mutants. The transcript was downregulated in the treated wild type compared to the control (FC of −3.7 WT NF/WT) and upregulated in the treated *gun1* and *gun5* mutant compared to the treated wild type with a FC of 3.9 and 4.2, respectively. This lncRNA transcript is able to overlap with an mRNA encoding an UDP-Glucose transferase (AT1G05560). Furthermore, like the lncRNA the overlapping mRNA transcript was downregulated in the treated wild type (FC of −5.8 WT NF/WT) and upregulated in both treated mutants (FC of 3.6 *gun1* NF/WT NF and 3.1 *gun5* NF/WT NF). We also detected differentially expressed sRNA clusters that can be processed from this region in the sRNA sequencing data in the treated wild type compared to the control (FC of −4.4 WT NF/WT) as well as in the treated *gun1* mutant (FC of 6.6 *gun1* NF/WT NF). Thus, the regulation of the nat-siRNAs is in agreement with the expression changes of the underlying lncRNA/mRNA transcript pair and given the differences between wild type and *gun* mutants the differential expression seems to be regulated by specific retrograde signalling pathways. In addition, we identified a *TAS3* precursor transcript (AT3G17185) that was downregulated in the NF treated wild type (−2.9 WT NF/WT) and was de-repressed in the treated *gun5* (2.6 *gun5* NF/WT NF). Ta-siRNAs produced from the *TAS3* transcript control the expression of transcripts coding for auxin response factors such as ARF2, ARF 4 and ETT. However, we neither detected differentially expressed *TAS3*-derived ta-siRNAs nor differential expression of their cognate targets between the analysed samples.

### Differentially regulated nuclear and organellar encoded mRNAs after NF treatment

In parallel to sRNA and lncRNA we analysed the data obtained from the ribosomal depleted nuclear (Figure 7) and organellar (Figure 8) encoded RNA sequencing to identify protein-coding mRNAs that are regulated by retrograde signalling pathways. In both untreated *gun1* and *gun5* mutants we detected only a low number of DEGs compared to the untreated wild type (Figure 7a) with 212 and 165 differentially expressed transcripts in the untreated *gun1* (*gun1*/WT) and *gun5* (*gun5*/WT) mutant, respectively. However, when we analysed differential gene expression in response to NF we observed a remarkable increase in the number of DEGs (Figure 7b). We identified 1557 transcripts that were differentially regulated in the wild type in response to NF as compared to the untreated control (WT NF/WT) including 412 upregulated and 1145 downregulated transcripts. For both treated mutants compared to their respective untreated controls we identified slightly lower numbers of DEGs. In total, 1361 DEGs were identified in the treated *gun1* mutant compared to the untreated *gun1* and 486 DEGs were upregulated and 875 DEGs downregulated. In the treated *gun5* 1177 DEGs were detected where 515 DEGs are up- and 662 DEGs are downregulated compared to the untreated *gun5*. In addition, we compared mRNA expression between the NF treated *gun* mutants and the NF treated wild type. We identified 905 DEGs in the NF treated *gun1* mutant compared to the treated wild type (587 DEGs upregulated and 318 DEGs downregulated) and 1319 DEGs in the treated *gun5* mutant (980 DEGs upregulated and 339 DEGs downregulated). We generated a hierarchically clustered heatmap from all 3352 nuclear encoded mRNAs that were differentially regulated in at least one sample (Figure 7c). Based on the co-expression of DEGs we were able to separate 15 specific clusters of differentially regulated nuclear encoded genes (Table S7). We identified many differentially expressed genes (1557 DEGs) in the treated wild type compared to the untreated control where nearly 75% of the mRNAs are downregulated. As expected the NF treated *gun* mutants behave in an opposite manner compared to the treated wild type since the majority of the RNAs were upregulated with 65% and 75% upregulated DEGs in the treated *gun1* (*gun1* NF/WT NF) and *gun 5* mutant (*gun5* NF/WT NF), respectively.

**Figure 7.**
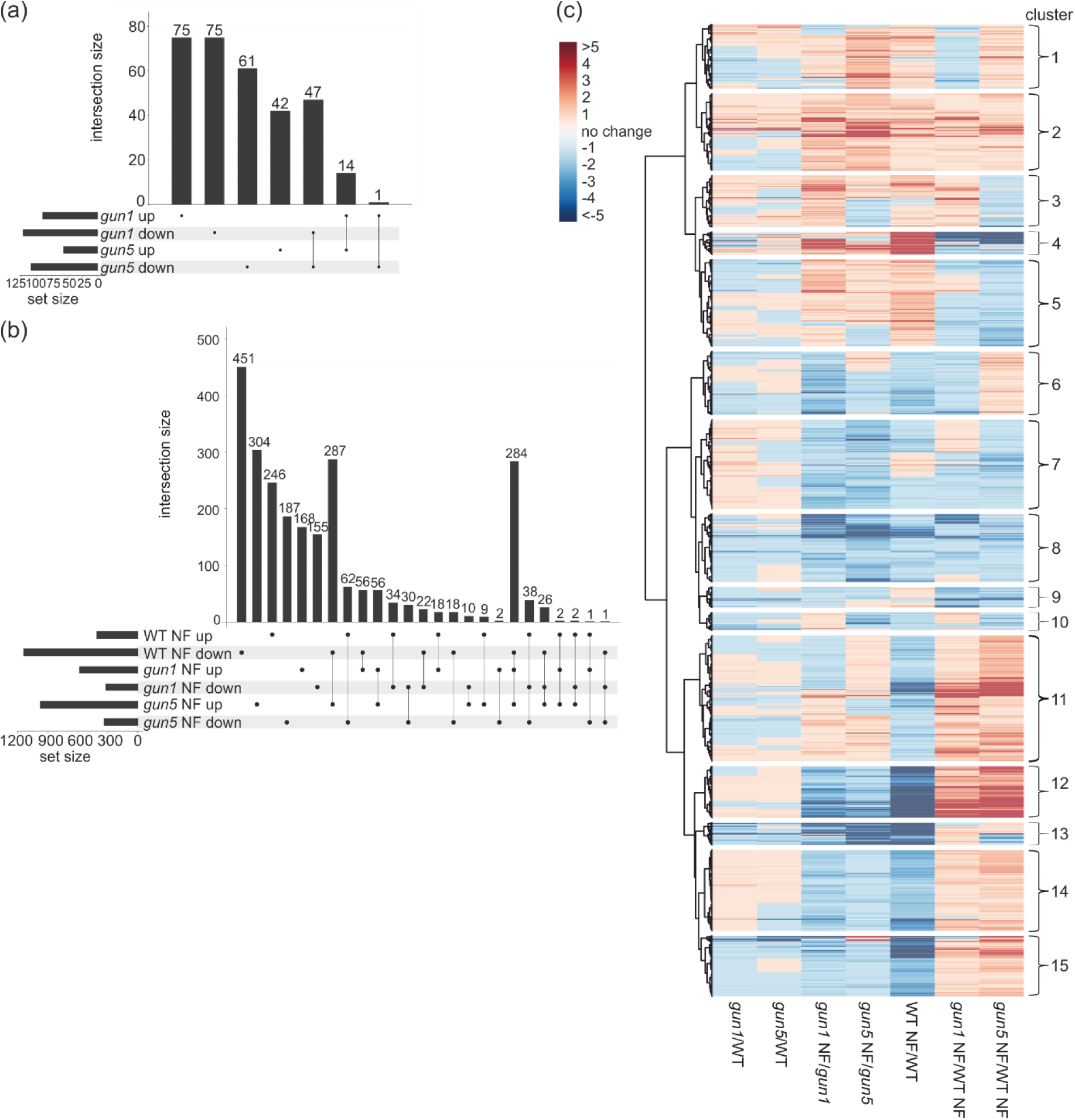
Distribution of differentially expressed nuclear DEGs in the untreated and NF treated samples. (a) UpSet plot showing the distribution of differentially regulated mRNAs (FC ≤-2 and ≥+2; FDR ≤ 0.05) in the untreated *gun* mutants compared to the WT. (b) UpSet plot depicting the distribution of differentially regulated mRNAs in response to NF in WT (WT NF/WT) and both *gun* mutants (*gun1* NF/WT NF and *gun5* NF/WT NF). (c) Hierarchically clustered (UPGMA) heatmap depicting nuclear encoded differentially expressed genes with 15 clusters based on co-expression patterns.

**Figure 8.**
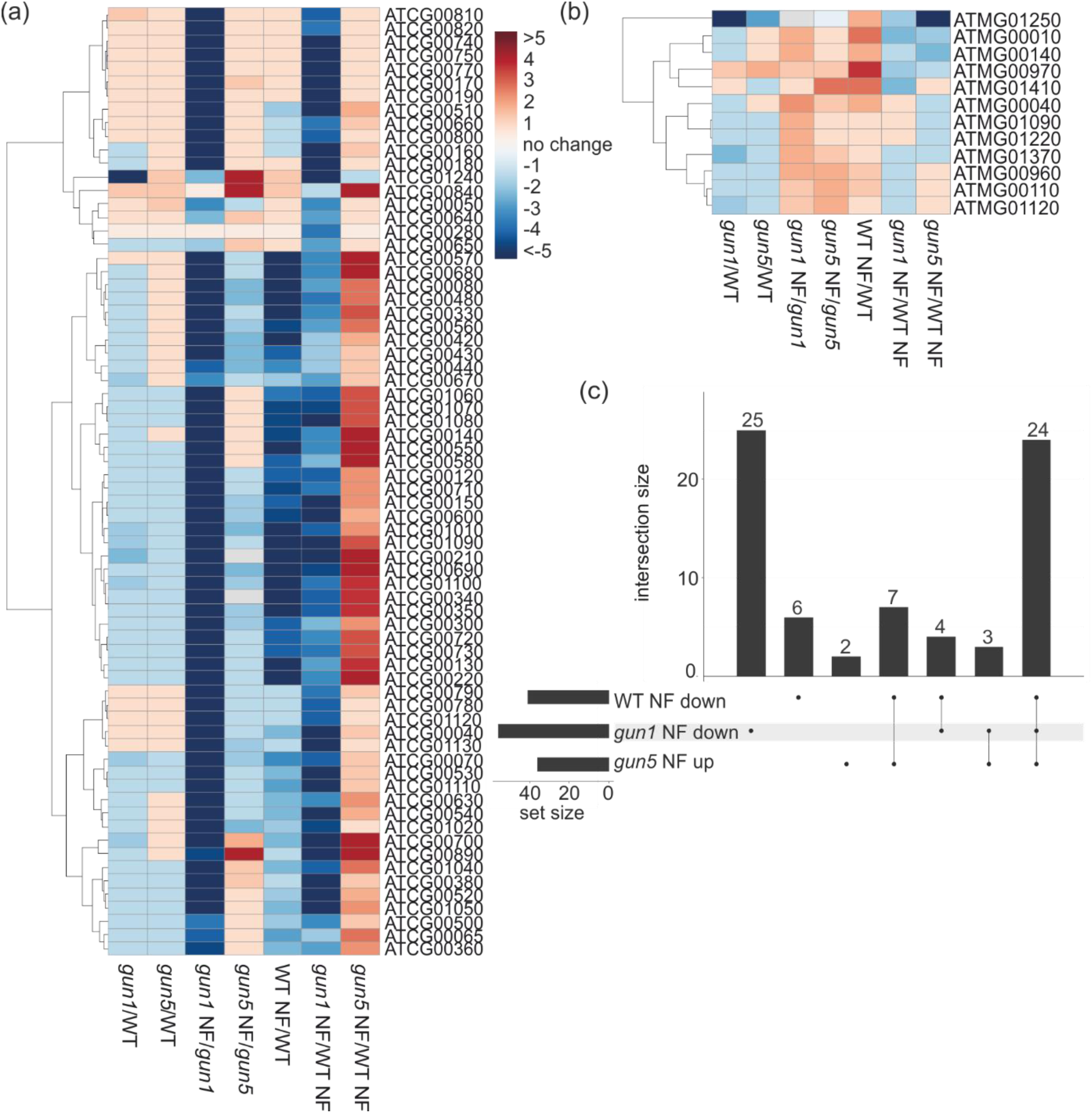
Distribution of differentially expressed plastidic and mitochondrial DEGs in the untreated and NF treated samples Hierarchically clustered (UPGMA) heatmap depicting plastidic (a) and mitochondrial (b) genes that are differentially expressed (FC ≤-2 and ≥+2; FDR ≤ 0.05) in at least one of the samples. (c) UpSet plot depicting the expression of plastid encoded DEGs detected in the NF treated *gun* mutants (*gun1* NF/WT NF and *gun5* NF/WT NF) and in the NF treated WT (WT NF/WT).

To identify the most interesting candidates that are regulated by retrograde signals we analysed the overlap between the treated wild type (WT NF/WT) and both treated *gun* mutants (*gun1* NF/WT NF and *gun5* NF/WT NF) to detect those genes that display a typical *gun*-related expression in both mutants (Figure 7b). We identified 284 DEGs which were differentially regulated in response to NF in wild type compared to the untreated control as well as differentially regulated in response to NF in both *gun* mutants compared to the NF treated wild type. These DEGs seem to be controlled by retrograde signalling pathways, since they are repressed by NF in the wild type and de-repressed in the *gun* mutants. Furthermore, we detected 56 DEGs with a specific de-repression in the treated *gun1* mutant (*gun1* NF/WT NF) and another 287 DEGs specifically de-repressed in *gun5* (*gun5* NF/WT NF). Most likely the regulation of the genes requires specific retrograde signals as we identified genes that show a specific de-repression that is restricted to only one of the *gun* mutants.

Besides the analysis of nuclear encoded genes, we investigated organellar gene expression and studied the expression of genes encoded by the plastidic and mitochondrial genome in the wild type and both *gun* mutants in the absence or presence of NF. We generated two hierarchically clustered heatmaps for plastidic (Figure 8a) and mitochondrial (Figure 8b) genes that were differentially expressed in at least one of the samples. As expected we only detected eight mitochondrial genes with differential expression level in at least one of the samples since NF treatment affects carotenoid biosynthesis in the plastids and should not directly affect mitochondrial gene expression. Furthermore, the low number of affected genes in the mitochondria indicates an insignificant cross-talk between plastids and mitochondria triggered by plastid-derived retrograde signals.

In contrast to the low number of differentially expressed mitochondrial genes we detected a considerable high number of differentially regulated plastid encoded genes. Upon growth in the absence of NF none of the plastid encoded genes was differentially expressed in the *gun5* mutant and only one plastid encoded gene was differentially expressed in the *gun1* mutant as compared to the untreated wild type. However, after NF treatment we detected 41 differentially expressed plastid encoded genes in the treated wild type compared to the untreated control and 56 and 36 DEGs in the treated *gun1* and *gun5* mutant, respectively, compared to the treated wild type. Furthermore, we noticed a highly interesting phenomenon with respect to the differentially expressed plastid encoded genes in both *gun* mutants. Whereas almost all plastid encoded differentially expressed mRNAs were downregulated in the NF treated *gun1* mutant compared to the treated wild type (*gun1* NF/WT NF) all plastid encoded differentially expressed mRNAs in the NF treated *gun5* mutant were upregulated as compared to the treated wild type (*gun5* NF/WT NF). Thus, based on the plastidal gene expression both mutants respond in an almost completely opposed manner to the NF treatment suggesting specific perturbations in the NF-triggered organellar signalling pathways. We observed 27 plastid encoded mRNAs that were differentially expressed in response to NF in both the *gun1* and *gun5* mutant (Figure 8c) when compared to the NF treated wild type, but were regulated in an opposing manner in the mutants: in the treated *gun1* mutant all were downregulated while they were upregulated in the treated *gun5* mutant.

#### Gene Ontology terms enrichment (mRNA)

To categorise putative functions of differentially regulated RNAs after NF treatment, Gene Ontology (GO) enrichment terms were explored and we considered GO-term enrichments that passed the statistic test with FDR ≤ 0.05 (Table S8 and Figure S2). In total, we obtained 24 significant enriched GO-terms for biological processes, 10 for cellular compartments and seven for molecular functions.

In the category of biological processes, we found considerable GO term enrichments in the treated wild type compared to the untreated control. The most abundant GO terms were reductive pentose-phosphate cycle (GO:0019253), positive regulation of catalytic activity (GO:0043085) and cellular response to sulfur starvation (GO:0010438). This indicates that NF treatment of wild type affects different processes like the Benson-Calvin cycle which takes place in the plastid and metabolic disturbances associated with catalytic proteins. In both NF treated *gun* mutants the reductive pentose-phosphate cycle (GO:0019253), protein-chromophore linkage (GO:0018298) and photosynthesis (GO:0015979) were highly enriched pointing to an overrepresentation of molecular processes and biochemical pathways that take place within the plastid. Moreover, we also noticed a strong enrichment of GO terms related to cellular compartments that are located in the plastid. The NAD(P)H dehydrogenase complex (plastoquinone) (GO:0010598), light-harvesting complex (GO:0030076) and stromule (GO:0010319), which all reside in the plastid are the most enriched GO terms in the treated wild type as well as in the treated *gun* mutants. These analyses imply that changes in the processes taking place in the plastid are the key targets in response to NF treatment. In the molecular function category, the enriched GO terms upon NF treatment in wild type and both *gun* mutants also include plastid-associated functions like pigment binding (GO:0031409), chlorophyll binding (GO:0016168) and poly(U) RNA binding (GO:0008266) with enriched plastid RNA binding proteins. In summary we conclude that processes mainly occurring in the plastids are key targets effected in the *gun* mutants with NF treatments.

### miRNA target analysis

miRNAs are able to regulate transcript levels by binding to reverse complementary RNA targets to mediate their cleavage. To correlate the expression of miRNAs with putative target RNA transcripts, we performed target prediction using all protein coding and non-coding transcripts from Araport11 (https://apps.araport.org/thalemine/dataCategories.do). For this, we used the sRNA target prediction tool “psRNATarget” and carried out miRNA target prediction with an expectation value ≤ 2.5 and RNA accessibility with a maximum UPE (unpair energy) of 25 (Table S9). For each predicted miRNA target we have considered its expression changes to subclassify the miRNA:RNA pairs. For the differentially regulated miRNAs, which were detected in WT NF/WT, *gun1* NF/WT NF and *gun5* NF/WT NF we were able to predict 218 protein-coding targets as well as 16 non-coding target RNAs and some of these can be targeted by several miRNAs. We generated a non-redundant list of miRNA targets and excluded transcripts with low FPKM values (≥ 5). Applying these parameters, we obtained 119 predicted miRNA targets that were further categorised into three different classes based on their expression and it has to be noted that a specific miRNA:RNA pair can be grouped into different categories since the miRNA as well as the cognate RNA target can be differently regulated between the analysed samples. The first category comprises miRNA:RNA pairs that are “unchanged” according to the fold change of the RNA transcript (but the miRNA does) and those transcripts with a q-value of more than 0.05, which include 101 miRNA:RNA pairs. The second category contains seven miRNA:RNA pairs that show an anticorrelated expression pattern and the third category encompasses 16 miRNA:RNA pairs where the miRNA as well as the predicted target show the same direction of their differential expression (both up- or both downregulated). Scatter plots (Figure 9) were created to show the distribution of the differentially regulated miRNAs and their correlating targets and were divided into “direct” and “indirect” scatter plots. The direct plots show the correlation of miRNAs and their cognate RNA targets either coding for transcription factors or coding for other proteins. The indirect scatter plots depict the expression of downstream genes that are controlled by miRNA-regulated transcription factors. From the direct scatter plot it is obvious that most differentially expressed miRNA targets do not encode transcription factors. Nevertheless, we identified transcription factor transcripts which are controlled by miRNAs and their effect on the transcription factor targets can be seen in the indirect plots. In the NF treated wild type compared to the untreated control the indirect plot shows many differentially expressed transcripts coding for transcription factors which are controlled by miRNAs. This effect cannot be seen in both treated *gun* mutants compared to the untreated control.

**Figure 9.**
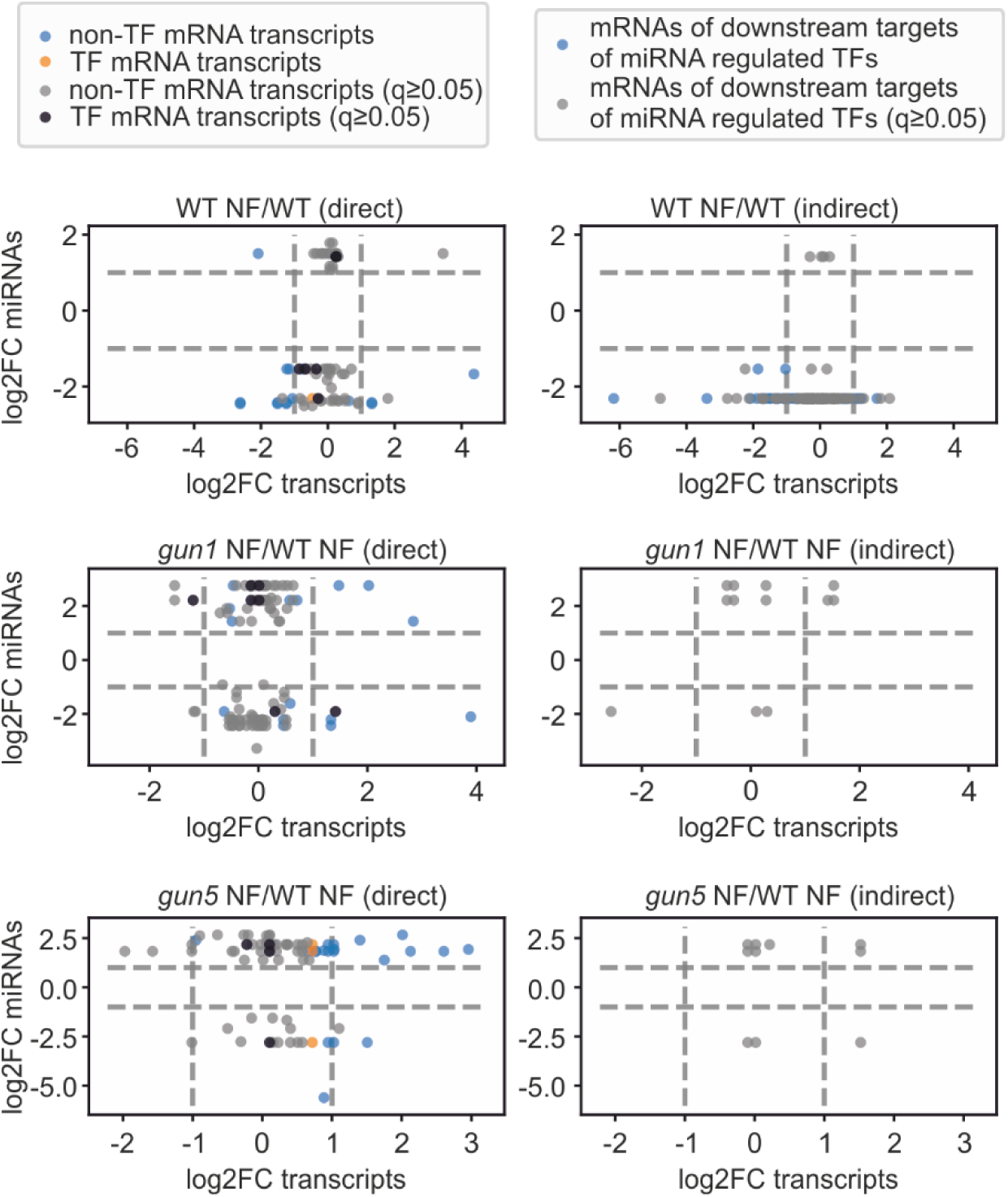
Scatter plots of differentially expressed miRNAs and their targets. Only miRNAs with FDR ≤ 0.05 were included. The direct plots (left panel) depict all differentially expressed miRNAs and their direct predicted target transcripts. miRNA target transcripts encoding transcription factors are shown in orange (FDR ≤ 0.05) and black (FDR ≥ 0.05). miRNA target transcripts encoding other proteins are shown in blue (FDR ≤ 0.05) and grey (FDR ≥ 0.05). The indirect plots (right panel) depict downstream targets of transcription factors that are miRNA-regulated. Here, the mRNAs of these downstream genes are plotted against the miRNAs controlling their respective transcription factor mRNAs. The blue dots correspond to a FDR ≤ 0.05 and the grey dots to a FDR ≥ 0.05.

We identified one miRNA:RNA target pair (Table S9) that has been shown to play a role in the acclimation to phosphate deficiency. miR399a is downregulated in the treated *gun1* mutant compared to the NF treated wild type whereas the expression of its target *PHO2* (AT2G33770) encoding an ubiquitin-conjugating E2 enzyme remains unchanged in the treated *gun1* mutant compared to the NF treated wild type. miR850 and its cognate target (AT1G09340) encoding a chloroplast RNA binding protein belong to the category of miRNA-target pairs showing the same expression (Table S9) since both are upregulated in the treated *gun5* mutant compared to the NF treated wild type. This chloroplast RNA binding protein is necessary for the proper function of the chloroplast and mutations in this gene cause growth deficiency (Fettke et al., 2011). Furthermore, we also identified a miRNA:RNA pair (Table S9) displaying an anticorrelated expression. miR157a-5p is strongly downregulated (−7 FC) in the treated *gun5* mutant compared to the NF treated wild type whereas its target AT4G28660 (PHOTOSYSTEM II REACTION CENTER PSB28 PROTEIN) is 2.9 fold upregulated (*gun5* NF/WT NF). PSB28 is highly conserved in photosynthetic eukaryotes and lack of *PSB28* results in a pale-green phenotype in rice pointing to a role in the assembly of chlorophyll containing proteins such as *CP47* (Lu, 2016).

### nat-siRNA target analysis

We detected a larger number of differentially expressed sRNAs arising from predicted NATs-pairs than from any other sRNA class in the treated wild type compared to the untreated control as well as in both NF treated *gun* mutants compared to the treated wild type. Filtering the differentially expressed nat-siRNAs with at least five normalised reads in all six samples (WT, WT NF, *gun1*, *gun1* NF, *gun5*, *gun5* NF) led to a total number of 73 non-redundant *cis*-NATs and 193 non-redundant *trans*-NATs pairs. These pairs were further analysed and we only selected the underlying nat-siRNA producing transcript pairs with at least five normalised reads for one of the two overlapping transcripts that reduced the number to 64 non-redundant *cis*-NATs and 40 non-redundant *trans*-NATs pairs (Table S10).

For many *trans*-NATs pairs we observed that one of the transcripts is derived from a transposable element or a pre-tRNA whereas the second overlapping transcript presents a protein coding gene. Among these *trans*-NATs pairs we only detected one overlapping transcript that encodes a plastid localised protein suggesting a low impact of *trans*-NATs pairs in the adjustment of plastid and nuclear gene expression in response to NF. The *trans*-nat-siRNA generated from this pair was found to be downregulated in the treated wild type compared to the untreated control and upregulated in both treated *gun* mutants compared to the treated wild type. The first overlapping transcript codes for the plastid localised UDP-glucosyl transferase 75B2 (AT1G05530), which is able to bind UDP-glucose important for cellulose and callose synthesis (Hong et al., 2001). Its expression was unchanged in the treated wild type compared to the untreated control as well as in both treated *gun* mutants compared to the treated wild type. The second overlapping transcript presenting an lncRNA (AT1G05562) was downregulated in the treated wild type compared to the untreated control and upregulated in both treated *gun* mutants compared to the treated wild type.

Interestingly out of 64 *cis*-NATs pairs that give rise to differentially regulated nat-siRNAs we detected 31 individual transcripts which encode plastid proteins indicating a considerable role of *cis*-NATs pairs in the direct control of genes coding for plastid proteins via NF triggered retrograde signals. Moreover, within the *cis*-NATs pairs we identified 35 individual transcripts encoding nuclear localised proteins pointing to a large impact of these in the indirect adjustment of nuclear gene expression via nuclear regulatory proteins. One sRNA processed from a *cis*-NATs-pair was detected to be downregulated in the treated wild type after NF compared to the untreated control. Interestingly, both overlapping transcripts were identified to encode plastid localised proteins. The expression for the first overlapping transcript (AT1G29900) is unchanged and it codes for a subunit of carbamoyl phosphate synthetase, which is presumed to be necessary for the conversion of ornithine to citrulline in the arginine biosynthesis pathway (Molla-Morales et al., 2011). In agreement with the expression of the nat-siRNAs the second overlapping transcript (AT1G29910) coding for a plastid localised protein is downregulated by NF in the wild type. This transcript encodes a chlorophyll A/B binding protein which is the major protein of the light-harvesting complex and required for absorbing light during photosynthesis.

### Network analysis

In order to gain a comprehensive picture of the role of miRNAs in retrograde signaling and to analyse possible downstream effects, we investigated a miRNA:RNA-target network that also comprises related transcription factor to target gene connections. Combining both results in a complex interaction network (Data S1), since one miRNA can control many mRNAs encoding transcription factors which in turn control several downstream genes, but also one miRNA target can be controlled by numerous miRNAs (Figure 10, Table S12). Within the considered network most miRNAs regulate just a small number of target transcripts (Figure 10a), but there are some miRNAs that are able to regulate up to 140 targets. In contrast, the majority of miRNA targets are regulated by only a few miRNAs, but there are still some targets that can be regulated by up to 15 miRNAs (Figure 10b). We observed that those miRNAs controlling the highest number of targets mainly regulate mRNAs that do not encode transcription factors (Figure 10c), while the distribution of miRNA targets encoding transcription factors, indicates that most miRNAs regulate only a small number of such targets with the highest number being 8 (Figure 10d). Some motifs are recurrent in the miRNA:RNA-target network (Figure 11). We explored the network for different characteristic relations of regulatory linkage and behaviour. Here we found simple, expected patterns where a miRNA is downregulated and its target mRNA in turn is upregulated or vice versa (Figure 11a). But we also observed many miRNA targets that did not show any differential expression on the mRNA level, although corresponding miRNAs were differentially expressed. The effect of these miRNAs might be visible on the protein level due to inhibition of translation. If the target mRNA encodes a transcription factor, we should then see the miRNA dependent regulation in the expression of downstream targets of this transcription factor (Figure 11b), as has been reported before (Megraw et al., 2016). As an example, the transcription factor AT4G36920 can act as an activator or repressor on its targets. If we consider it acts as an activator and the miRNA controlling this transcription factor mRNA is downregulated, we would expect an upregulation of the transcription factor mRNA and, hence, an upregulation of its downstream target transcripts. In case of repression, the transcripts of corresponding downstream targets should be downregulated, because of enhanced repression. This is not the case for its downstream target AT3G16770, which is upregulated in the NF-treated wild type compared to the untreated control. Also, the considered transcription factor acts as an activator on AT1G14350, but the corresponding mRNA is downregulated in the treated wild type compared to the untreated control, which is contrary to our expectations. Besides mRNAs of transcription factors, many miRNAs are able to regulate transcripts of genes that do not encode transcription factors, but also these transcripts do not always show the expected behaviour (Figure 11c). Further, we find examples for a single downstream target that can be controlled by several miRNAs (Figure 11d). In these cases, behavioural predictions are impossible without knowing more about the exact mode of action of each miRNA and the magnitude of their influence.

**Figure 10.**
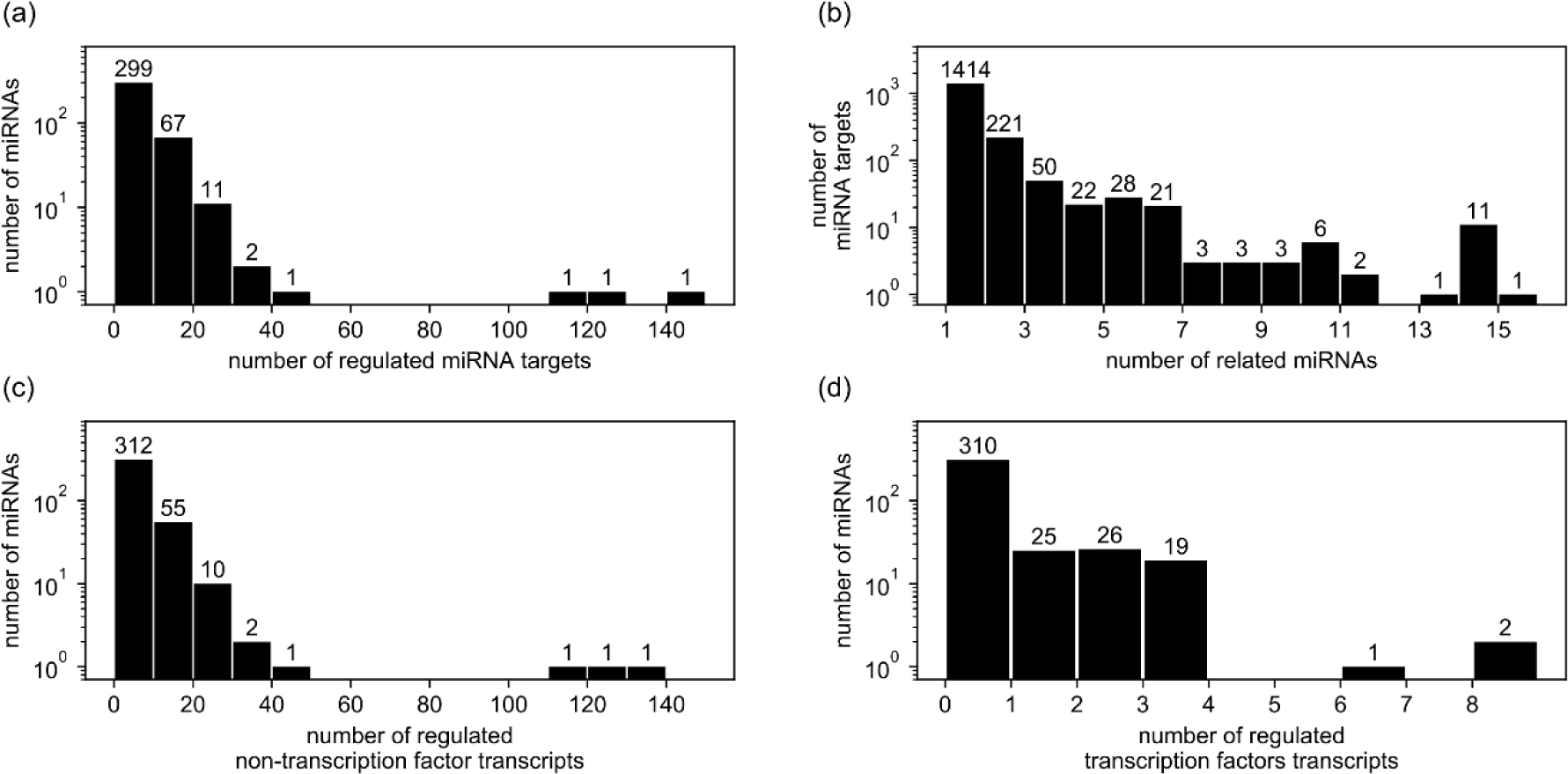
Distributions of miRNA:RNA-target interaction numbers (Table S12). The first distribution (a) shows the amount of miRNAs regulating a certain number of putative RNA-targets. The second distribution (b) shows the amount of predicted RNA-targets that are regulated by a certain number of miRNAs. The bottom two distributions again show the amount of miRNAs regulating a certain number of putative RNA-targets separated by RNA-targets encoding non-transcription factors (c) or transcription factors (d). For (a) and (c) the data was binned in groups of 10. The number on top of each bar indicates the number of members of each bin.

**Figure 11.**
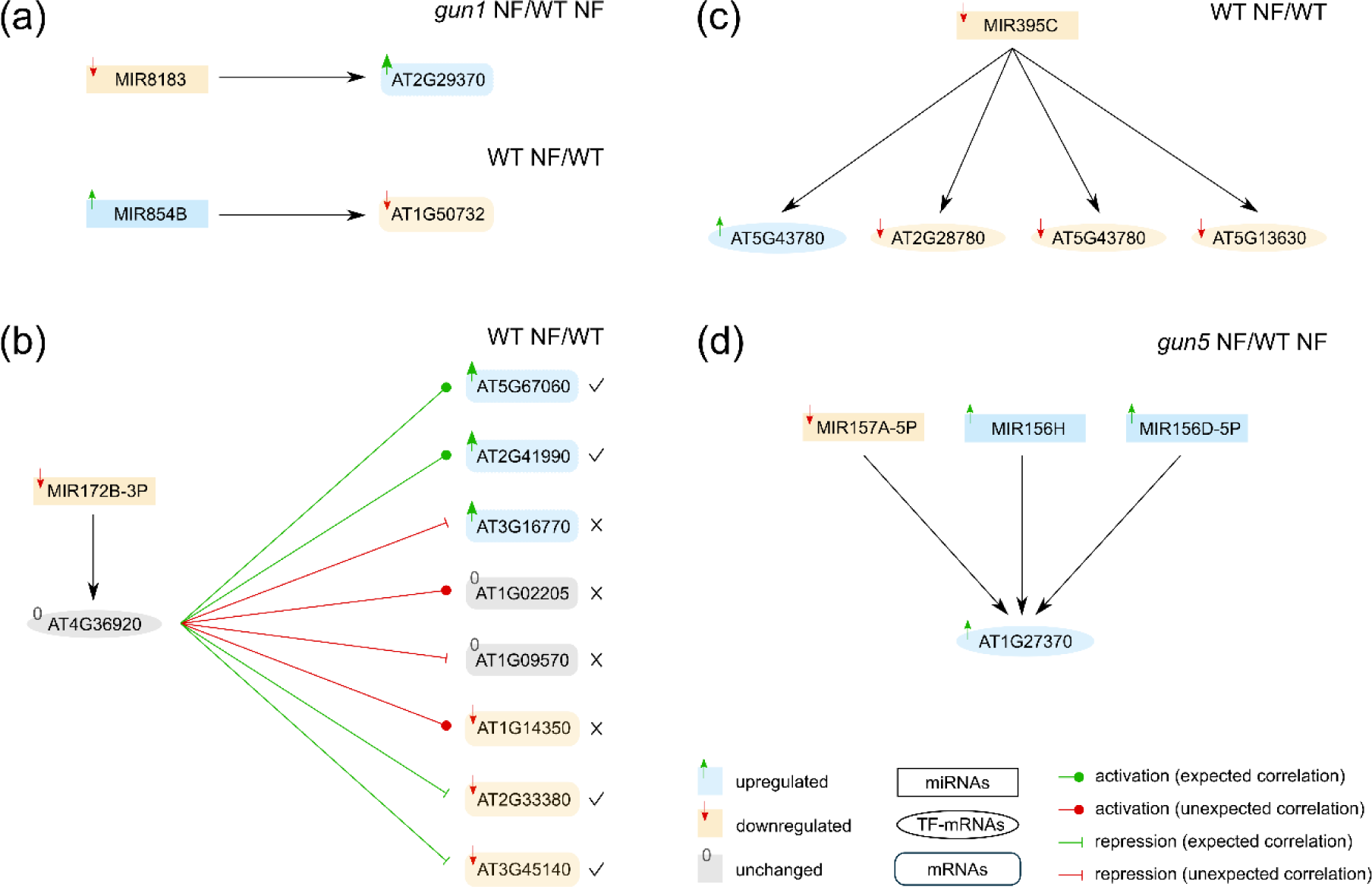
Illustration of different network motifs which we observed in the miRNA:RNA-target network in connection with relative changes of RNA levels between treatments. The first scheme (a) show examples for expected regulations where a miRNA and its target mRNA exhibit inversed differential expression. Scheme (b) shows the interaction between a miRNA that regulates the mRNA of a transcription factor coding gene, and the interactions of this transcription factor with its downstream target genes. The labelled shapes represent transcripts of the corresponding genes. Their colours reflect relative changes between the indicated measurements (upper right). Cases were classified as “expected correlation” if the change in expression of the mRNA matched the expected behaviour, taking into account the regulatory effects of miRNA and transcription factor. Otherwise it was classified as “unexpected correlation”. The classification for each case is indicated by the colour of the interaction marker and the checkmarks or crosses (expected correlation: green and checkmark; unexpected correlation: red and cross). While the miRNA is downregulated in the given case, the transcription factor mRNA is not differentially expressed and downstream targets of this transcription factor exhibit varying mRNA expression patterns. Scheme (c) shows a downregulated miRNA which regulates four different targets. Scheme (d) shows an example of three miRNAs that regulate a single mRNA.

## Discussion

Until now nothing is known whether ncRNAs and sRNAs are regulated by retrograde signalling in response to norflurazon (NF) treatment and how they contribute to the control of nuclear gene expression in response to plastid-derived signals. To understand these concepts, we combined sRNA sequencing with mRNA/lncRNA sequencing of *A. thaliana* wild type seedlings as well as seedlings of the two retrograde signalling mutants *gun1* and *gun5* which were treated for four days with the herbicide norflurazon. Both *gun* mutants have defects in the retrograde signalling pathway which lead to a de-repression of the *PhANGs* after NF treatment and therefore are suitable to identify ncRNAs and mRNAs which are regulated by retrograde signals.

Generally, after NF treatment we detected nearly the same number of DEGs in all treated samples compared to the untreated wild type. Further, we observed a general tendency that more DEGs were downregulated than upregulated in response to the NF treatment. In addition, we could observe an overrepresentation of differentially expressed genes encoding plastid localised proteins in all three samples and detected more DEGs to be upregulated in the treated *gun* mutants compared to the treated wild type.

Previous studies with different *gun* mutants were performed using microarrays representing *A. thaliana* protein coding genes, but lacking probes for ncRNAs to detect retrograde-dependent changes in gene expression (Strand et al., 2003, Koussevitzky et al., 2007). A study by Koussevitzky *et al*. (2007) was based on a very similar experimental setup as used in our study where the authors analysed changes in mRNA levels in wild type (*Col-0*), *gun1* and *gun5* mutant seedlings that were grown for five days on media with and without NF and we compared the results from this study with our data (Figure S3). Koussevitzky *et al*. (2007) identified 2587 DEGs (1038 upregulated and 1549 downregulated) in the NF treated wild type compared to the untreated control and thus detected 1644 more DEGs (up- and downregulated) in response to NF compared to our results (Figure S3a and b). Furthermore, 177 upregulated DEGs and 766 downregulated DEGs were commonly detected in both studies and both studies identified a larger number of downregulated DEGs and larger changes in the NF treated *gun5* mutant compared to the treated wild type than in the treated *gun1* mutant. There is also a good overlap of DEGs that were found in both studies in *gun1* and *gun5* mutants. About 56 % of the DEGs detected in the *gun1-102* (*gun1* NF/WT NF) in our study were also detected the treated *gun1-9* mutant by Koussevitzky *et al*. (2007) (*gun1* NF/WT NF) (Figure S3c) and about 50 % of the DEGs identified in the treated *gun5* in our study were also identified by in Koussevitzky *et al*. (2007) (*gun5* NF/WT NF) (Figure S3d). However, in our data set we also identified 44 % (*gun1* NF/WT NF) and 50 % (*gun5* NF/WT NF) of DEGs that have not been shown to be controlled by *gun*-related retrograde signalling pathways before which might be due to the differences between both methods (RNAseq vs. microarrays) and the varying duration of the NF treatment in the studies (five days vs. four days).

In contrast to the differentially expressed nucleus encoded genes which show large overlaps between the *gun1* and *gun5* mutant we observed an opposite regulation of differentially expressed plastid encoded transcripts in both *gun* mutants. In response to NF all differentially expressed plastid encoded transcripts were downregulated in the *gun1* mutant whereas they were upregulated in the treated *gun5* mutant. These observations are in line with the model suggesting that plastid gene transcription is controlled by retrograde signalling networks including sigma factors (SIG2, SIG6) and plastid encoded RNA polymerase (PEP) which might be crucial for proper plastid RNA transcription (Woodson et al., 2013). It seems that GUN1 activates PEP (Maruta et al., 2015) and a perturbed PEP activation in the *gun1* mutant may prevent the upregulation of the plastid encoded genes in the treated *gun1* mutant compared to the treated wild type.

We identified an interesting scenario where one lncRNA (AT4G13495) was found to be differentially regulated in our lncRNA sequencing data showing the classical de-repression in both *gun* mutants compared to the treated wild type. This lncRNA overlaps in sense direction with three different miRNA precursors (MIR5026, MIR850 and MIR863). In our sRNA sequencing data sRNAs that can be derived from this lncRNA and all three miRNAs were also detected to be differentially expressed in at least one treatment (WT NF/ WT, *gun1* NF/WT NF and *gun5* NF/WT NF) and we assume that all three miRNAs can be produced either from the three individual miRNA precursor transcripts or from the lncRNA. We did not predict any target for miR5026 according to the applied psRNATarget parameters. miR850 is upregulated in the *gun5* mutant (*gun5* NF/WT NF) and two predicted cognate target RNAs encoding a chloroplast RNA binding protein and a threonine-tRNA ligase, respectively, were upregulated as well. miR863 is targeting the *SERRATE* transcript (AT2G27100) encoding an accessory protein that is essential for the miRNA biogenesis pathway and thus may influence the regulation of several miRNAs (Meng et al., 2012). miR863 did not show a significant differential expression in the treated wild type (WT NF/WT), but it was upregulated in both treated *gun* mutants compared to the treated wild type. However, we did not detect significant changes of the *SERRATE* transcript in both treated *gun* mutants.

Concerning the overall regulation of differentially expressed sRNAs belonging to different sRNA classes (miRNAs, nat-siRNAs and other sRNA producing loci) we detected more downregulated sRNAs in the treated wild type compared to the untreated control (WT NF/WT) whereas both treated *gun* mutants exhibit a higher number of upregulated sRNAs compared to the treated WT (*gun1* NF/WT NF and *gun5* NF/WT NF). Principally, this suggests an increased sRNA processing in response to NF in both *gun* mutants resembling the de-repression of nuclear encoded genes and we assume that these sRNAs may have an impact on retrograde-controlled nuclear gene expression. sRNAs are able to affect nuclear transcripts that are regulated by retrograde signals and they may regulate mRNA transcripts that might affect plastid localised proteins. In all sRNA classes we observed the highest number of differentially regulated sRNAs within the nat-siRNA class in all treatments.

We could identify the maximum downregulation with respect to the expression level in the treated wild type compared to the untreated control for the miRNA miR169g-3p (−151.6 fold WT NF/WT). The same miRNA showed the highest detected upregulation with a value of 38.2 fold in the treated *gun5* mutant compared to the treated wild type (*gun5* NF/WT NF). We did not predict any target for miR169g-3p according to our parameters. Generally, miR169 has already been shown to act in stress-responses including heat and salt stress (Szyrajew et al., 2017, Pegler et al., 2019) and regulates targets such as the *NUCLEAR FACTOR Y, SUBUNIT A5* that was shown to be important in the heat stress response. In tomato leaves miR169g-3p was reported to be downregulated under Pi deficiency, but a target for this particular miRNA could not be predicted (Gu et al., 2010).

We found 119 non-redundant predicted miRNA targets for 47 differentially regulated miRNAs in the treated wild type (WT NF/WT) as well as in both treated *gun* mutants (*gun* NF/WT NF). In detail, 101 predicted miRNA targets belong to the category “unchanged”, seven targets belong to the class of “anticorrelated” targets and 16 targets belong to the group showing the “same correlation”. The expression of most of the targets is not anticorrelated to the expression changes of their cognate miRNA which leads us to conclude that miRNAs either do not have such drastic effects on the expression of genes that are controlled by retrograde signalling pathways or the expressional changes of miRNAs somehow balance transcriptional changes of their targets to maintain constant steady-state levels or they act as translational repressors and do not have a direct effect on the transcript abundance of their target RNAs. We predicted 20 miRNA targets that code for transcription factors and 22 targets encoding plastid localised proteins which are targeted by differentially regulated miRNAs. Thus, we assume that miRNAs may have important functions in the control of transcripts that code for regulatory proteins that are directly involved in transcriptional control and may contribute to the manifold changes of gene expression in response to retrograde signals. Further, nuclear transcripts that code for plastid localised proteins like the chloroplast RNA binding protein (AT1G09340) are targets for miRNAs and suggest that these specific miRNA:mRNA pairs can play an important role in the retrograde signalling pathway, because the miRNA directly controls transcripts coding for plastid proteins and thus may contribute to the adjustment of plastidal and nuclear gene expression. One interesting case involves miR395b and miR395c which target the mRNA for the magnesium-chelatase subunit (AT5G13630). In the NF treated wild type both miRNAs and the target mRNA are downregulated compared to the untreated control whereas in the treated *gun5* mutant both miRNAs as well as the target are upregulated. Thus, the expression of this miRNA:mRNA pair is not anticorrelated, but might still be interesting, since the enhanced miRNA levels may balance an increased transcription rate of the target mRNA to keep physiologically relevant steady-state levels. The magnesium-chelatase is required in the chlorophyll biosynthesis pathway where it catalyses the insertion of Mg^2+^ into protoporphyrin IX and the *gun5* mutant is characterised by a single nucleotide substitution causing a defective magnesium-chelatase. In the wild type the *GUN5* transcript decreases in response to NF triggered the retrograde signalling whereas the transcript level in the *gun5* mutant lacking the specific retrograde signal remains high and cannot be efficiently downregulated by the increased miRNA levels. The detected classical anticorrelation of the miRNA:mRNA pairs point to regulatory functions of specific miRNAs in the retrograde signalling pathway since we assume efficient miRNA-mediated target cleavage followed by a reduced mRNA steady-state level. In this category of anticorrelated pairs we identified the mRNA for the *SPL10* transcription factor representing a validated target of miR157a suggesting that miR157a acts in retrograde signalling by affecting the levels of a transcriptional regulator and its downstream targets. Another anticorrelated predicted miRNA:mRNA pair is miR398 targeting the transcript of the MATE (multidrug and toxic compound extrusion) efflux protein (AT2G04050). We detected miR398 to be downregulated in the NF treated wild type compared to the untreated control and the target is slightly upregulated. This MATE efflux protein belongs to a huge class of membrane proteins which can be located in the plasma membrane and the chloroplast envelope membrane (Wang et al., 2016). These proteins are able to bind cytotoxic compounds like primary and secondary metabolites and eliminate them from the cell (Liu et al., 2016). They also transport metabolic and xenobiotic organic cations like norfloxacin and ethidium (Omote et al., 2006) and additional toxic substances such as pollutants, herbicides or xenobiotic products out of the cell (Diener et al., 2001). miR398 was downregulated in response to NF most likely causing elevated transcript levels and increased levels of the encoded plasma membrane located MATE efflux protein. It can be speculated that the regulated MATE efflux protein might be involved in the extrusion of the applied herbicide NF or the extrusion of toxic compounds that accumulate within the cell response to NF treatment.

Beside this, we also identified a miRNA target encoding for a plastid protein that appeared to be downregulated in the NF treated *gun5* mutant (*gun5* NF/WT NF) by increased levels of the cognate miRNAs. *PSB28* (AT4G28660) is targeted by miR157a and encodes a protein that is part of the photosystem II reaction centre and is suggested to function in the biogenesis and assembly of chlorophyll-containing proteins. miR157a is downregulated in the NF treated *gun5* mutant whereas the *PSB28* mRNA shows an anticorrelated upregulation (*gun5* NF/WT NF). NF treatment usually leads to the downregulation of *PhANGs* and thus should cause decreased expression levels of the *PSB28* mRNA. However, miR157a that may contribute to the downregulation of *PSB28* mRNA levels posttranscriptionally seems to be controlled by retrograde signals indicated by the misregulation of miR157a in the *gun5* mutant. Thus, we assume that the increased *PSB28* transcript levels most likely are caused by the misregulation of miR157a that shows reduced levels in the *gun5* mutant.

Fang et al. (2018) reported that tocopherols are required for the accumulation of 3′-phosphoadenosine 5′-phosphate (PAP) presenting a retrograde inhibitor of the nuclear exoribonucleases (XRN) which may protect primary miRNAs from being degraded and promote mature miRNA production. In addition, the transcript encoding the enzyme ATP sulfurylase (*APS*) that catalyses the initial step in PAP synthesis (Klein and Papenbrock, 2004) was identified to be targeted by miR395 (Liang et al., 2010). Fang et al. (2018) reported that PAP is involved in the regulation of miRNA biogenesis via the proposed inhibition of XRNs since they detected reduced PAP levels and concomitant increased levels of pri-miRNAs and mature miRNAs in *MIR395f* overexpression lines. We detected this interesting miRNA:mRNA pair as differentially expressed our sRNA sequencing data. miR395b was downregulated in the treated wild type (WT NF/WT) as well as upregulated in the treated *gun5* mutant (*gun5* NF/WT NF). Further, we also detected the *APS* mRNA as a target of miR395 to be upregulated in the treated wild type (WT NF/WT) and unchanged in the treated *gun5* mutant (*gun5* NF/WT NF). In the wild type NF causes reduced miR395b levels leading to an increase of the *APS* transcript most likely causing elevated PAP synthesis that acts as retrograde inhibitor of XRNs and should provoke elevated pri-miRNA and mature miRNA levels. Beside this Fang et al. (2018) detected the downregulation of miR398 in the wild type after NF treatment which is in line with our our sRNA sequencing data since we also identified miR398 to be downregulated in the treated wild type (WT NF/WT). The COPPER/ZINC SUPEROXID DISMUTASE2 (*CSD2*) was previously detected to be the target of miR398 (Guan et al., 2013). After heat stress Fang, et al. (2018) detected miR398, PAP and tocopherols to be increased and *CSD2* to be decreased in the wild type and they hypothesise that tocopherols and PAP are required for miR398 biogenesis under heat stress. The *CSD2* mRNA escaped our miRNA target prediction due to a considerably high number of mismatches within the miRNA binding site causing a score value that was above our cut-off value. Still we identified this miRNA as differentially expressed supporting the previous study by Fang et al. (2018). Generally, these two miRNAs which were identified by Fang et al. (2018) to be important in the retrograde signalling could also be confirmed in our study applying NF as another trigger of retrograde signalling and we identified additional miRNAs and their putative targets which seem to have an regulatory function in the retrograde signalling pathway.

Beside the differentially expressed miRNAs we identified an even higher number of differentially regulated nat-siRNAs in the treated wild type compared to the control as well as in the treated *gun1* and *gun5* mutant compared to the treated wild type. Most of the overlapping transcripts encode for nuclear or plastid proteins suggesting that nat-siRNAs predominantly control transcripts encoding transcription factors that confer retrograde controlled gene expression and that nat-siRNAs have a considerable impact on the control of *PhANGs* encoding plastid proteins. We identified one *cis*-NATs-pair which showed a classical anticorrelation with regard to RNA transcript and sRNA expression level. One of the overlapping transcripts (AT1G05562) belonging to this *cis*-NATs-pair was identified to be differentially expressed in the lncRNA data set (upregulated in WT NF/WT and downregulated in *gun1* NF/WT NF and *gun5* NF/WT NF) and the related nat-siRNA was determined to be downregulated in the treated wild type compared to the control as well as upregulated in the treated *gun1* and *gun5* mutant compared to the treated wild type. The gene AT1G05562 encodes an antisense lncRNA and overlaps with the gene AT1G05560, which codes for an UDP-glucose transferase. Both, the transcript and the nat-siRNA show anticorrelated expression levels. After NF treatment the nat-siRNA increases which then causes a decrease of one of the overlapping transcripts in the wild type whereas in both *gun* mutants the nat-siRNA decreases and causes increased transcript levels. Another interesting *cis*-nat-siRNA pair was identified to be downregulated in the treated wild type (WT NF/WT) and upregulated in the treated *gun5* mutant (*gun5* NF/WT NF). The first overlapping gene (AT1G29930) encoding a chlorophyll binding protein was downregulated in the treated wild type (WT NF/WT) and upregulated in the treated *gun5* mutant (*gun5* NF/WT NF) whereas the overlapping transcript coding for a nuclear RNA polymerase (AT1G29940) remained unchanged in both treatments (WT NF/WT and *gun5* NF/WT NF). Considering the regulation of the expression levels, the results suggest that in case of the treated *gun5* mutant the nat-siRNA is controlling the transcript of the chlorophyll binding protein, since an increase in nat-siRNAs leads to a decrease in the transcript upon NF triggered retrograde signalling. Here, we could demonstrate that NF treatment and subsequent retrograde signals lead to comprehensive changes in the steady-state levels of non-coding sRNAs comprising all known sRNA classes. The majority of the identified differentially expressed sRNA belong to the *cis*- and *trans*-nat-siRNAs followed by the second most abundant class presenting miRNAs. Thus, we postulate that mainly these two sRNA classes act as important regulators of gene expression in retrograde signalling. We also identified a considerable high number of so far unknown differentially expressed nuclear-encoded genes and thus add to the knowledge about genes that are controlled by retrograde signalling. Finally, we were able to detect promising sRNA-RNA target pairs that may act in the adjustment of plastidic and nuclear gene expression in retrograde signalling pathways.

## Experimental procedures

### Plant material and growth conditions

*Arabidopsis thaliana* wild type (*Col-0*) and the retrograde signalling mutants *gun1-102* and *gun5-1* were used in this study. *gun1-102* (SAIL_290_D09) harbours a T-DNA insertion within the AT2G31400 gene locus resulting in a loss-of-function allele (Tadini et al., 2016). *gun5-1* is an EMS mutant harbouring a point mutation within the gene AT5G13630 causing an Ala → Val substitution at residue 990 (A990V) resulting in a deficient magnesium-protoporphyrin IX synthesis (Mochizuki et al., 2001). Surface-sterilised seeds were incubated on ½ MS agar plates containing 1.5% sucrose. For treatments with norflurazon (NF), seeds were incubated on the same medium supplemented with 5 µM norflurazon (Sigma-Aldrich). After vernalisation (2 days at 4°C in darkness) the seeds were grown for four days under continuous light (115 µmol photons m^−2^s^−1^) at 22°C. Whole plants were harvested and immediately frozen in liquid nitrogen and stored at −80°C until RNA isolation. All control experiments and norflurazon treatments were performed in three biological replicates for each genotype.

### RNA isolation

The plant material was ground in liquid nitrogen and RNA isolation was performed using Trizol reagent (Invitrogen) according to the manufacturer’s protocol. RNA integrity was monitored by agarose gel electrophoresis and RNA concentration and purity was determined spectrophotometrically (260 nm/280 nm and 260 nm/230 nm absorbance ratios).

### sRNA purification

For sRNA sequencing 30 µg of total RNA were loaded on a 15% PAGE and separated for 2 h at 120V. The sRNA fractions with sizes ranging from 18 to 29 nucleotides were excised from the gel and eluted in 0.3 M NaCl overnight at 4°C with rotation. Remaining gel pieces were removed using a Spin-X centrifuge tube (Sigma-Aldrich) and 1 µl GlycoBlue (15 mg/ml, Thermo Fisher), 25 µl sodium acetate (3 M, pH 5.0) and 625 µl ethanol were added to the 250 µl flow through and incubated for 4 h at −80°C. After centrifugation for 30 min with 17.000 × g at 4°C the RNA pellet was washed twice with 80% ethanol, dried and dissolved in nuclease free water.

### Quantitative RT-PCR

Prior to cDNA synthesis RNA samples were subjected to DNase I digestion (NEB) to remove residual genomic DNA. 2 µg total RNA was incubated at 37°C for 30 min together with DNaseI (2 U, NEB). To inactivate the DNaseI, 2.5 µl 50 mM EDTA were added and incubated at 65°C for 10 min. The RNA was denatured for 5 min at 65°C in the presence of 100 pmol of an oligo-dT23VN oligonucleotide and 10 mM dNTPs and transferred to ice. Subsequently, cDNA synthesis was performed for 1 h at 42°C using M-MuLV reverse transcriptase (200 U, NEB) followed by a heat-inactivation at 80°C for 5 min. To monitor successful cDNA synthesis, we performed reverse transcription (RT) PCR using gene-specific primers for the gene *UBI7* (AT3G53090) (Table S11).

For each qRT-PCR we used cDNA equivalent to 20 ng/µl RNA with gene-specific primers and an EvaGreen qPCR mix. The samples were pre-heated for 2 min at 95°C and qRT-PCR cycling conditions were: 12 sec at 95°C, 30 sec at 58°C and 15 sec at 72°C for 40 cycles. All qRT-PCRs were performed in three technical triplicates with the CFX Connect Real-Time PCR device (Bio-Rad). The ct-values were used to calculate changes in gene expression by ΔΔCt method (Livak and Schmittgen, 2001). The values were normalised to the housekeeping gene *UBI1* (AT4G36800). Oligonucleotide sequences of all gene-specific primers are listed in Table S11.

### RNA Sequencing

For the generation of mRNA libraries including polyA-tailed lncRNAs 10 µg total RNA from each sample were vacuum-dried in the presence of RNA stable (Sigma-Aldrich). The libraries were prepared using the Next Ultra RNA Library Prep Kit (NEB) by the company Novogene (China). The samples were sequenced strand-specifically as 150 bp paired-end on a HiSeq-PE150 platform with at least 15 million read pairs per library.

sRNA libraries for each RNA sample were generated twice following two slightly modified protocols. The first set of libraries was generated from 5 µg total RNA with the NEBNext Multiplex Small RNA Library Prep Set for Illumina according to the manufacturer’s instructions and 1 h of 3’ adapter ligation. The second set of sRNA libraries was prepared from purified sRNAs obtained from 30 µg of total RNA using the same Kit as described above performing 3’ adapter ligation for 18 h. For both libraries, excessive non-ligated 3’ adapters were made inaccessible by converting them into dsRNA by hybridisation of complementary oligonucleotides. 5’ adapter ligation was carried out at 25°C for 1.5 h, reverse transcription was performed by using the ProtoScript II reverse transcriptase and libraries were amplified by 12 PCR cycles. The PCR products were loaded on a 6 % PAGE and separated for 2 h at 60 V. The cDNA library fractions with a size ranging from 138 to 150 nucleotides were excised from the gel and eluted overnight. The sRNA libraries were sequenced as 50 bp single-end reads on an Illumina HiSeq1500 sequencer with approximately 10 million reads per library.

### Analysis of mRNA and lncRNA

The mRNA and lncRNA sequencing results were analysed with the open source and web-based platform GALAXY (Afgan et al., 2016). The FASTQ raw sequences were trimmed with the tool Trimmomatic to remove adapter sequences with their default parameters (Bolger et al., 2014). Tophat (Kim et al., 2013) was used to map the reads against the *A. thaliana* reference genome (https://www.arabidopsis.org/, release: TAIR10) with a maximum intron length parameter of 3,000 nt. The transcripts were annotated in Araport11 (https://apps.araport.org/thalemine/dataCategories.do) and besides mRNAs we considered annotated ncRNAs longer than 200 bp as lncRNAs. Differential expression of transcripts was analysed by Cuffdiff (Trapnell et al., 2010) to normalise the sequencing depth of each library and to calculate fragments per kilobase of transcript per million reads (FPKM). The false discovery rate (FDR) was used as a statistic indicator to exclude type I errors or rather false positives. Transcripts having FDR ≤ 0.05 and log2 fold change (FC) ≤-1 and ≥+1 were considered as differentially expressed genes (DEGs). The package pheatmap (https://cran.r-project.org/web/packages/pheatmap/pheatmap.pdf) was used to generate hierarchically (UPGMA) clustered heatmaps of differentially expressed RNAs (Kolde, 2019).

### Gene Ontology terms

Gene Ontology (GO) enrichment terms were analysed using the DAVID Bioinformatics Resources 6.8 (https://david.ncifcrf.gov) with default parameters (Huang da et al., 2009a, Huang da et al., 2009b) and results were visualised with the R package “ggpubr” (Wickham, 2016).

### Analysis of sRNA

NEBNext Kit adapter sequences were clipped from the sequencing reads using a custom script within GALAXY that identifies Illumina adapter sequences using a seed sequence of 10 nt. After adapter clipping FASTQ files of the raw reads with a length of 18 to 26 nt were loaded into the CLC Genomics Workbench 11.0.1 (Qiagen) for further analyses. The ShortStack analysis package was used for advanced analysis of the sRNA sequences (Axtell, 2013b). The FASTQ files of the six biological replicates derived from each treatment were first mapped against the *A. thaliana* TAIR10 reference genome (https://www.arabidopsis.org/, release: TAIR10). The merged alignments were mapped against a file covering all *A. thaliana* mature miRNAs (http://www.mirbase.org/) and a second file comprising different RNA classes, namely nat-siRNAs, ta-siRNAs, phased small interfering RNAs (phasiRNAs) and lncRNAs. A nat-siRNA database (Table S1) was generated from previously annotated NATs-pairs (Zhang et al., 2012, Yuan et al., 2015, Jin et al., 2008), phasiRNAs were taken from Howell *et al*. (Howell et al., 2007) and lncRNA were downloaded from Araport11 (https://apps.araport.org/thalemine/dataCategories.do, release: Araport 11 Annotation). After mapping to the respective references, the individual raw reads for each replicate were used for normalisation and differential expression analysis based on a calculation with DeSeq2 (Love et al., 2014). The sRNAs were filtered by fold changes between ≤ −2 and ≥ +2. The significance of the differentially expressed sRNAs was evaluated with q-value of ≤ 0.05. miRNA target RNAs were identified using the psRNATarget (Dai et al., 2018) prediction V2 tool (http://plantgrn.noble.org/psRNATarget/) from protein coding and non-coding transcripts present in Araport11. An expectation value of less than 2.5 was considered as a cut-off for true miRNA targets where mRNAs harbouring a lower number of mismatches to the reverse and complementary miRNA obtain lower score values.

### Network analysis

Using the sRNA and mRNA sequencing data together with the miRNA target prediction, we assembled an interaction network of miRNAs and their putative targets. For network analysis, we used the python package networkX (A. Hagberg et al., 2008). We further investigated the miRNA targets that were predicted as described above using the psRNATarget tool to identify all miRNA targets that encode transcription factors. For this, we compared all miRNA targets with a reference database, containing *A. thaliana* transcription factors, which was generated using publicly available data (http://atrm.cbi.pku.edu.cn). Furthermore, this reference database was extended by incorporating available information about whether the transcription factors act as activators or repressors of gene expression together with available information about the individual target genes of the transcription factors (https://agris-knowledgebase.org). The data obtained from the RNA sequencing experiments was then used to generate a network of miRNAs and their targets, differentiating between miRNA targets encoding transcription factors and targets encoding other proteins. For network analyses connected to measurements, only RNAs with a FDR less than 0.05 were considered, unless indicated otherwise. Network analyses were performed to determine relationships between miRNAs and their targets with special focus on miRNA targets encoding transcription factors. In these network analyses we considered the impact of miRNAs on the transcripts coding for transcription factors and the impact of the expression of transcription factor mRNAs on downstream genes, regulated by these transcription factors. If the change of a miRNA results in an expected change of an mRNA coding for a transcription factor, or at least downstream genes show expected transcriptional changes, we classify this behaviour as “expected”. For example, if miRNA expression is reduced, the target mRNA encoding a transcription factor is upregulated, the transcription factor acts as an activator and downstream genes of this transcription factor are consequently also upregulated, this would be considered as “expected” behaviour. Furthermore, the scatter plots presented for all differentially expressed miRNAs (FDR ≤ 0.05) were subdivided into two categories: plots depicting relations of miRNA and their direct target transcripts (direct) and plots depicting indirect relations comprising miRNAs that control mRNAs encoding transcription factors and their downstream target genes.

## Accession numbers

ATG accession numbers: GUN1 (AT2G31400), GUN5 (AT5G13630)

The raw Illumina sRNA and mRNA/lncRNA sequencing data is deposited in NCBI SRA database with the ID PRJNA557616.

## Acknowledgments

This work was supported by the German Research Foundation (SFB-TRR 175, grants to W.F. project C03 and E.K. project D03). We thank Martin Simon for technical support and advice regarding sRNA library preparations.

## Author contributions

WF and MAA designed the research; KH performed the research with help from MAA and BT; KH, MAA, BT and WF analysed the data; network analysis was performed by SOA, MK and EK; and KH, MAA, MK, SOA and WF wrote the paper. All authors read and approved the final manuscript.

## Short legends for Supporting Information

Figure S1. Validation of transcript levels by qRT-PCR.

Figure S2. Gene Ontology term enrichment analysis.

Figure S3. Comparison of a previous study to our study.

Table S1. Reference sequences for the *cis*- and *trans*-NATs-pairs.

Table S2. sRNA sequencing and mapping results for each independent biological replicate.

Table S3. Lists of all significant differentially expressed sRNA classes.

Table S4. mRNA mapping results after the tool Tophat for each independent biological replicate.

Table S5. Overview of all significant differentially expressed mRNAs.

Table S6. Lists of all significant differentially expressed mRNAs and other RNA classes.

Table S7. List of the individual clusters of the mRNA heatmap.

Table S8. Gene Ontology term enrichment analysis for significant DEGs.

Table S9. Lists of miRNA:RNA target prediction.

Table S10. Lists of *cis*- and *trans*-nat-siRNAs and their target correlation.

Table S11. Sequences of oligonucleotides used in this study.

Table S12. Distributions of miRNA:RNA-target interaction numbers.

Data S1. miRNA:RNA-target network in GML-format that can be opened with the free software “Gephi”(Bastian et al., 2009).

